# Integrative structural analysis of the human endosomal Na^+^/H^+^ exchanger NHE6 reveals a lipid-stabilized intracellular gate and a disordered C-terminus

**DOI:** 10.1101/2025.07.21.665523

**Authors:** Lukas P. Feilen, Emil E. Tranchant, Milena R. Lalic, Lara Sach, Jesper F. Havelund, Jan Ostendorf, Nils J. K. Færgeman, Stine F. Pedersen, Birthe B. Kragelund, Henriette E. Autzen

## Abstract

Human NHE6 (HsNHE6) is an endosomal Na^+^/H^+^ exchanger essential for maintaining luminal pH and endo-lysosomal trafficking in neurons. HsNHE6 mutations are implicated in devastating neurological syndromes, but mechanistically the transporter remains poorly understood. Here, we present the single particle cryo-electron microscopy structure of HsNHE6 at 3.4 Å, captured in an inward-open conformation. The structure reveals a homodimeric architecture with 13 transmembrane helices per protomer and the conserved ion-binding site. Functional assays demonstrate that HsNHE6 mediates the exchange of both Na^+^ and K^+^ for protons. A structured C-terminal helix interacts with the transmembrane core, jointly forming a hydrophobic cavity that accommodates two lipid molecules, potentially modulating cation access to the ion-binding site. The distal C-terminus of HsNHE6 is intrinsically disordered, as revealed by NMR and small-angle X-ray scattering and equips HsNHE6 with a large cytosolic span. Our integrative model of full-length HsNHE6 provides a comprehensive framework for understanding HsNHE6-mediated ion exchange and its disruption in neurological diseases such as Christianson syndrome.

## Introduction

Sodium/hydrogen exchangers (NHEs) are ancient proteins, highly conserved across all kingdoms of life^1^. As secondary active transporters, NHEs harness electrochemical gradients to exchange monovalent cations (Na^+^/Li^+^/K^+^) and protons (H^+^) across cellular membranes^2^. In mammals, NHEs are electroneutral and drive the exchange of monovalent cations, primarily Na^+^, for H^+^. These NHEs are encoded by the solute carrier 9 (SLC9) gene family, comprising 13 different isoforms divided into three sub-families: SLC9A (NHE1-9), SLC9B (NHA1-2), and SLC9C (SLC9C1 and SLC9C2)^3^. NHEs exhibit significant diversity in tissue distribution, subcellular localization, cation selectivity, and regulatory mechanisms^3^. Mammalian NHEs play critical roles in a wide range of physiological processes such as Na^+^ reabsorption in the kidney and intestine, tissue-, organellar and cytosolic acid-base homeostasis throughout the body, cell volume regulation in response to osmotic stress, and cellular processes like cell proliferation, migration, or vesicle trafficking^3^.

Although key aspects of NHE function remain unresolved, particularly the conformational states along the transport cycle, recent NHE structures provide a much-needed foundation for understanding their roles in disease and informing targeted therapeutic strategies. For instance, structural studies on bacterial and mammalian electroneutral NHEs, such as *Pyrococcus abyssi* NhaP (PaNhaP)^4^, *Methanocaldococcus janaschii* NhaP (MjNhaP)^5^, *Equus caballus* NHE9 (EcNHE9)^6^, as well as *Homo sapiens* NHE1^7^ and NHE3^8^ (HsNHE1 and HsNHE3) confirmed the homodimeric nature of NHEs inferred from early kinetic and biochemical studies^9–11^. Each NHE protomer consists of 12 or 13 transmembrane (TM) α-helices organized into a dimerization domain and a core domain, forming the ion translocation pore at their interface. The recent structures show that the ion translocation pore harbor highly conserved, negatively charged residues crucial for cation binding^3,7,8^. During the transport cycle these residues are alternately exposed to either side of the membrane, allowing for substrate engagement and release^4,5,7,12,13^. For NHEs, the substrate transport is hypothesized to happen through an “elevator” mechanism in which the core domain moves relative to the static dimerization domain, offering alternating access to the ion binding site^14^.

In contrast to the TM region, the cytoplasmic C-termini of NHEs are poorly conserved^3,15^ and contain a large fraction of structural disorder. Meanwhile, despite their variability in both length and sequence, the NHE C-termini are hypothesized to play important regulatory roles in localization and activity of NHEs and serve as hubs for protein and lipid binding partners^16–23^ as well as for regulatory phosphorylation and dephosphorylation events during which they may form defined, secondary structure elements^16,17^.

Within the NHE family, the isoforms NHE6, NHE7, and NHE9 are localized in intracellular membranes^3^. HsNHE6 (SLC9A6) is predominantly expressed in the nervous system where it mainly localizes to early and recycling endosomes^3,24,25^. Most of the available evidence suggests that HsNHE6 can exchange not only Na^+^, but also K^+^ for H^+^ contrasting plasma membrane-localized NHEs^26^. In the endosomal pathway, this enables HsNHE6-mediated transport to be driven by the prevailing endosomal Na^+^ and K^+^ gradients^27^ together with the H^+^ gradient generated by V-type ATPases^28^. As a result, HsNHE6 facilitates the endosomal exchange of Na^+^/K^+^ and H^+^, thereby playing a critical role in maintaining endosomal pH and regulating endo-lysosomal trafficking^29,30^.

Consistent with its key role in neuronal physiology, HsNHE6 dysfunction is implicated in severe neurological disorders, such as autism, Christianson syndrome (CS)^28,30–34^. Humans suffering from CS exhibit a range of mutations (truncations, deletions, point mutations) in HsNHE6. CS is one of the most prevalent X-linked neurodevelopmental disorders, affecting an estimated 1 in 16,000 to 1 in 100,000 individuals worldwide^31,35,36^. Intriguingly, CS presents a complex phenotype characterized by both neurodevelopmental and neurodegenerative effects, with symptoms that have similarities to those of lysosomal storage disorders^3^. Other neurodegenerative syndromes such as Alzheimer’s disease have been associated with reduced HsNHE6 expression in the brain^37,38^. The molecular understanding of how disease-causing mutations impact the structure and function of HsNHE6 has remained elusive, primarily due to the absence of a 3D structure and a full-length model.

To uncover the molecular basis of HsNHE6 function and its disruption in disease, we carried out an in-depth integrative structural and functional analysis of HsNHE6. Using single particle cryo-electron microscopy (cryo-EM), we determined the structure of HsNHE6 in an inward-open conformation, revealing a conserved ion-binding site and a structured C-terminal helix (C1) that interacts with the TM core. Lipidomics and structural data revealed co-purified lipids, including species bound near C1 that together with TM3 and TM4, form a hydrophobic cavity, where a conserved arginine appears to act as a cytoplasmic gate modulating access to the ion-binding site. Functional liposomal assays demonstrated that HsNHE6 mediates both Na^+^ and K^+^ exchange, with no detectable preference. Mass spectrometry identified a cleaved and acetylated N-terminus, and immunofluorescence microscopy (IFM) confirmed localization to early and recycling endosomes. Finally, NMR spectroscopy and small-angle X-ray scattering (SAXS) analyses showed that the distal C-terminus is intrinsically disordered, forming a dynamic extension likely involved in regulatory interactions. Together, these findings provide a full-length structural model of HsNHE6 and a mechanistic framework for understanding its role in ion homeostasis and its dysregulation in neurological disease.

## Results

### HsNHE6 has a dimeric architecture with 13 TM helices

To investigate the structure of HsNHE6, we expressed full-length HsNHE6 (Uniprot Q92581-2, residues: 1-701; **Supplementary Fig. 1**) with baculovirus transduction of FreeStyle 293-F cells (Hek293F; ThermoFisher Scientific) and purified it in the detergents glycodiosgenin (GDN) and cholesteryl hemisuccinate (CHS) (**Supplementary Fig. 2a**) for subsequent SP cryo-EM structure determination (**Supplementary Fig. 2b-d**). *Ab initio* reconstruction and three-dimensional (3D) classification produced four classes, one of which exhibited clear densities for TM α-helices. Subsequent 3D refinements applying C1 or C2 symmetry yielded cryo-EM maps with overall estimated resolutions of 3.5 Å and 3.4 Å, respectively, based on the Fourier shell correlation (FSC) gold-standard criterion at 0.143 (**Supplementary Fig. 3, Supplementary Table 1**). Although both cryo-EM structures were nearly identical, the structure with C1 symmetry revealed minor differences between the two protomers, particularly in the luminal loop 1 (LL1) between TM2 and TM3, which was better resolved in this structure (**Supplementary Fig. 4a**). Correspondingly, the two structural models show a root-mean-square-deviation (RMSD) of approximately 0.3 Å in their C^α^ atoms, indicating a high degree of structural similarity between them. (**Supplementary Fig. 4b**). LL1 extends over the dimerization interface, in close proximity to several densities interpreted as lipids in both the C1 and C2 cryo-EM reconstructions (**Supplementary Fig. 4c**). While neither the C1 nor the C2 cryo-EM maps allowed complete model building of LL1, we incorporated structural guidance from the EcNHE9 structural model^39^ and AlphaFold3^40^ predictions for the composite model (see later). Given the high similarity between the maps and the dimeric nature of HsNHE6, we focused the structural analysis on the C2-symmetrized model. The resolution of the TM helices and the remaining loops of the cryo-EM structure of HsNHE6 allowed automated *de novo* model building using ModelAngelo^41^ (**Supplementary Fig. 5**). HsNHE6 forms a dimer with a nearly rectangular cross-section when viewed both from the membrane plane and perpendicular to the membrane (**Fig. 1a**) like other eukaryotic NHEs^6–8^. Each protomer consists of 13 TM α-helices (TM1–TM13), arranged in a 6-TM structural-inverted repeat linked through a long α-helix, TM7: TM1-6 and TM8-13 exhibit a pseudo-2-fold symmetry along an axis parallel to the membrane plane, characteristic of transporters with the NhaA fold^2^. Within each protomer, TM1-3 and TM8-10 form the dimerization domain, providing the oligomerization interface for the two protomers. For each protomer, TM4-6 and TM11-13 form the ion transporting (core) domain (**Fig. 1b**). The N-terminus of HsNHE6 is positioned on the luminal side and the C-terminus on the cytoplasmic side of the membrane; However, the cryo-EM structure is missing parts of both the N-terminus and the C-terminus, respectively (**Fig. 1b**).

**Fig. 1.**
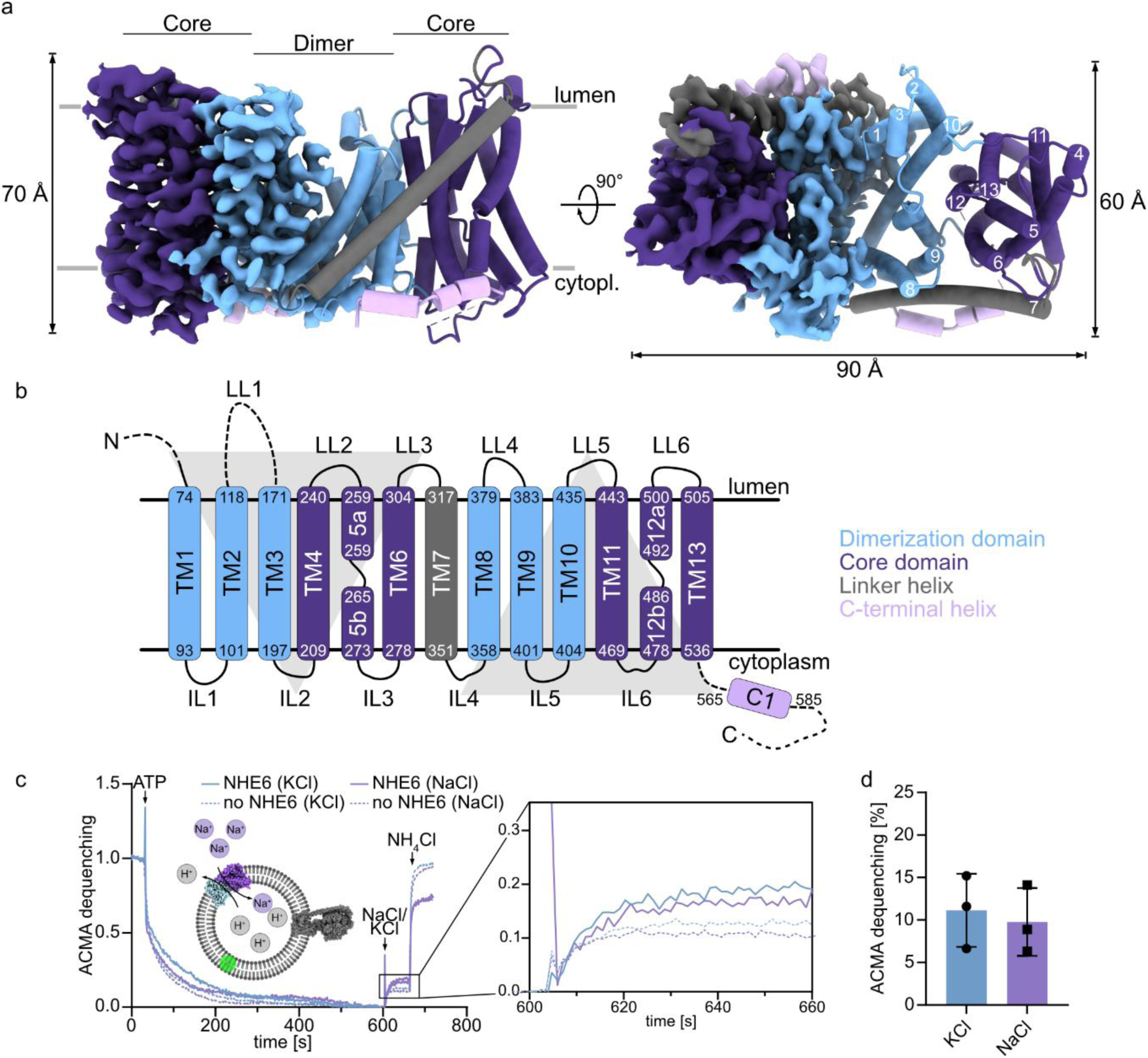
Architecture and activity of HsNHE6. **a,** The 3D structure of HsNHE6 TM domain represented from the side (left) and top (right) relative to the membrane plane. One protomer is depicted as the cryo-EM reconstruction and the other as the corresponding structural model. The colors represent the domain organization within each protomer, which is composed of the dimer domain (light blue, TM1-3 and TM8-10), the core domain (dark purple, TM4-6 and TM11-13), the linker helix (gray, TM7) and the C-terminal helix (C1, light purple). Cryo-EM densities that could not be attributed to protein are omitted for clarity. **b,** Schematic representation of the helix organization within one protomer of HsNHE6 colored as in a. The dotted lines represent parts that are not resolved in the cryo-EM structure. The numbers represent the TM boundaries, while luminal loops and intracellular loops are abbreviated LL and IL, respectively. **c,** Activity of HsNHE6 in Brain Polar Lipid Extrac (BrainPLEx) lipids containing CHS. Representative fluorescence traces of liposomes containing HsNHE6 and ATPase (solid lines) or only ATPase (dashed lines), respectively. Transport activity is induced by the addition of 150 mM KCl (violet traces) or NaCl (green traces), respectively. **d**, De-quenching of ACMA (9-Amino-6-Chloro-2-Methoxyacridine) after the addition of 150 mM KCl (violet) or NaCl (green), after baseline subtraction (mean ± SD, n = 3 biological replicates).

### HsNHE6 contains an N-terminal signal peptide and is localized in endosomes

At the outset of this study, it was unclear whether HsNHE6 comprised an uneven or even number of TM helices, a distinction with direct implications for the topology of the protein and the orientation of its termini. With 13 TM helices in the cryo-EM structure, the N- and C-termini are placed on opposite sides of the membrane. The absence of the first 73 N-terminal amino acid residues in the cryo-EM structure may be attributed to flexibility and/or the presence of an N-terminal signal peptide that may regulate the localization of HsNHE6^42^. While the SignalP 6.0 server^43^ predicts cleavage of a signal peptide just before S43 (**Supplementary Fig. 6a**), peptide mapping using MS analysis of trypsin-digested HsNHE6, identified a semi-tryptic peptide beginning at L32 as the most N-terminal peptide in the mature HsNHE6 (**Supplementary Fig. 6b**). These results are broadly consistent with a previous study proposing residues P27-F37 to form a short α-helix that targets HsNHE6 to the ER and is subsequently cleaved^42^. In addition to mapping how HsNHE6 is proteolytically cleaved during maturation, LC-MS/MS was also used to identify that the N-terminus is acetylated, preventing N-terminal sequencing by Edman degradation.

Given the post-translational modification of the N-terminus, features that are often implicated in protein trafficking and localization, we next sought to determine the subcellular distribution of HsNHE6. For this, FreeStyle293-F cells were transiently transfected with a pcDNA3.1(+) expression vector encoding HsNHE6-FLAG. Following fixation, cells were processed for IFM analysis to assess co-localization of the FLAG epitope with markers for the plasma membrane and specific cellular compartments (**Supplementary Fig. 7**). Confocal imaging revealed prominent localization of HsNHE6-FLAG with Rab5-positive early endosomes and transferrin receptor-positive early/recycling endosomes. Substantial localization of HsNHE6-FLAG at the plasma membrane was also observed. HsNHE6-FLAG co-localized to some extent with LAMP-1-positive lysosomes and partially with Giantin-positive Golgi structures. No appreciable co-localization was observed with endoplasmic reticulum and mitochondria. These observations are consistent with the reported localization pattern of HsNHE6 in cultured cells^29,44^.

### HsNHE6 transports K^+^ and Na^+^ in exchange for H^+^

We next assessed whether detergent-purified HsNHE6 retains ion exchange activity and whether it shows a preference for Na^+^ or K^+^ as the transported cation *in vitro*. Due to the localization of HsNHE6 in endosomes, we hypothesized that it could transport K^+^ as well as Na^+^ as previously proposed based on analysis of survival of salt-sensitive yeast overexpressing HsNHE6 in Na^+^ and K^+^ rich medium^26^. For this, HsNHE6 was reconstituted in liposomes composed of Brain Polar Lipid Extract (BrainPLEx) and CHS together with the F_0_F_1_-ATPase from *E. coli*. H^+^ transport, initiated by addition of either Na^+^ or K^+^, was monitored through the (de-)quenching of the proton-gradient-sensitive fluorophore ACMA^45^ (9-Amino-6-Chloro-2-Methoxyacridine, **Fig. 1c**). Upon the addition of ATP, ACMA was quenched, followed by de-quenching after the addition of either KCl or NaCl (**Fig. 1c**), showing that H^+^ flux by HsNHE6 can be induced by both K^+^ and Na^+^. The dequenching signal was around two-fold higher than the signal from BrainPLEx liposomes containing only the *E. coli* F_0_F_1_-ATPase. There was no detectable difference in the dequenching induced after addition of 150 mM KCl or 150 mM NaCl, respectively (**Fig. 1d**), indicating that under the conditions tested, HsNHE6 is equally capable of transporting K^+^ and Na^+^ against H^+^, without detectable preference.

### HsNHE6 adopts an inward-facing conformation with the ion binding site occupied by additional cryo-EM density

In the HsNHE6 structure, the core domain adopts an inward-facing conformation, as recognized by a funnel-shaped cavity connecting the ion-binding site located between the dimerization and core domains to the cytoplasmic space (**Fig. 2a**). The surface of this funnel is predominantly negatively charged (**Fig. 2a**), presumably facilitating the binding and transport of Na^+^, K^+^ and H^+^. Furthermore, TM5 and TM12, both part of the core domain, are unwound at their crossover point as observed for other transporters with an NhaA fold^12,46^. Comparison of the HsNHE6 model with published structures of HsNHE1, HsNHE3, and EcNHE9 reveal notable structural differences. The RMSDs of C^α^ atoms between HsNHE6 and HsNHE1 or HsNHE3 are approximately 2 Å, with significant variations observed in the loop regions and between TM1 and TM2 (**Supplementary Fig. 8a, b**). In contrast, the RMSD between HsNHE6 and EcNHE9 is around 1 Å, with differences primarily observed in the loops (**Supplementary Fig. 8c**).

**Fig. 2.**
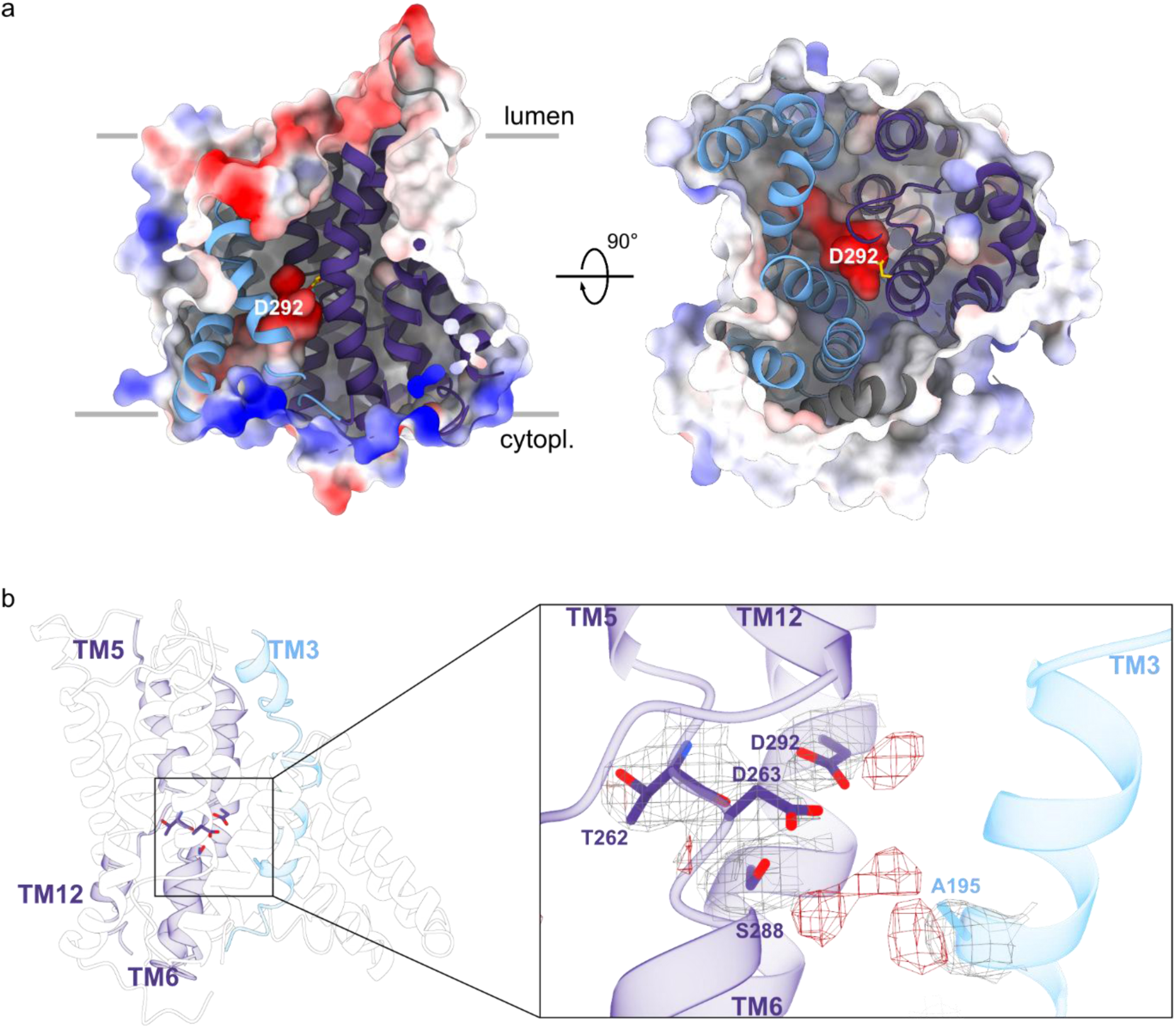
The ion binding site of HsNHE6. **a,** Electrostatic surface representation and structural model of one HsNHE6 protomer, viewed in the membrane plane (left) and from the cytosolic side (right) with the funnel between the dimerization (blue) and core domains (violet). The highly conserved ion-binding residue D292 is depicted in yellow sticks. **b,** Cartoon representation of the putative ion binding site between TM3, -5, -6, and -12 (colored by their respective domain), with an enlarged view onto the putative ion coordinating residues A195, T262, D263, S288, and D292 overlayed with the cryo-EM map around these residues. The red regions in the cryo-EM map correspond to unassigned densities in the ion binding site.

A previous study has shown that NHEs in the inward-facing conformation can accommodate a monovalent cation in their ion-binding site^5^. In the HsNHE6 cryo-EM structure, we observe unassigned densities near the residues of the ion-binding site composed of side chains of D263^TM5^, S288^TM6^, and D292^TM6^ as well as the backbone oxygen of T262^TM5^ (**Fig. 2b**). Here superscripts denote the TM α-helix to which each residue belongs). While the resolution of the HsNHE6 cryo-EM structure is insufficient for unambiguous density assignment, we hypothesize that the multiple densities in the ion binding site reflect varying Na+ occupancy across inward-open particles, rather than simultaneous binding of multiple ions (**Fig. 2b**). D292^TM6^ is highly conserved among NHEs, and the corresponding residues were found to be directly involved in ion transport in HsNHE1^47^, and NhaA^12^ and NapA^48^ from *Escherichia coli*. Furthermore, the arrangement of the side chains of the residues of the proposed ion binding site in the HsNHE6 structure closely resembles those of homologous residues in the crystal structure of Tl^+^-bound PaNhaP^4^ (PDB: 4CZA; **Supplementary Fig. 9a**). The ion binding site of PaNhaP has an additional negatively charged residue, E73^TM3^, which in the crystal structure of PaNhaP coordinates the Tl^+^ ion; however, this glutamic acid is not necessary for transport activity^4^. Interestingly, this glutamic acid is missing in all mammalian PaNhaP homologues, including HsNHE6. Beyond the additional densities, the arrangement of the ion binding site of HsNHE6 is analogous to that observed in the homologous HsNHE1 structure (**Supplementary Fig. 9b**), corroborating the conserved structural features of ion transport among human NHEs. Additionally, the residues E287^TM6^ and R457^TM11^ in HsNHE6 form a salt bridge, likely stabilizing the ion binding site (**Supplementary Fig. 10a**), which is further stabilized by hydrogen bond interactions between N291^TM6^ and T262^TM5^ and T219^TM3^, respectively. This E^TM6^-R^TM11^ salt bridge has previously been suggested to stabilize the ion binding site of NHEs^49^ and was also recently observed for EcNHE9^39^ (**Supplementary Fig. 10b**).

### Lipid molecules may regulate a cytoplasmic gate in the vicinity of a C-terminal α-helix

The C-terminus has remained largely unresolved in previously published NHE cryo-EM structures, likely due to their intrinsic disorder^3^. Notably, 20 amino acid residues of the C-terminus in HsNHE6 were resolved as an α-helix (W565–S585, designated C1). C1 is positioned on the cytosolic side of HsNHE6 and lies in the membrane plane near TM4, - 7, and -9, bridging the core- and dimerization domains (**Fig. 3a**). At this position, residues in C1 directly interacts with the highly conserved K350^TM7^ through hydrogen bonds with the backbone carbonyls of L581^C1^ and L582^C1^ similar to what was recently described for a cysteine-engineered variant of EcNHE9 (EcNHE9*CC) designed to stabilize the inward open conformation^39^.

**Fig. 3.**
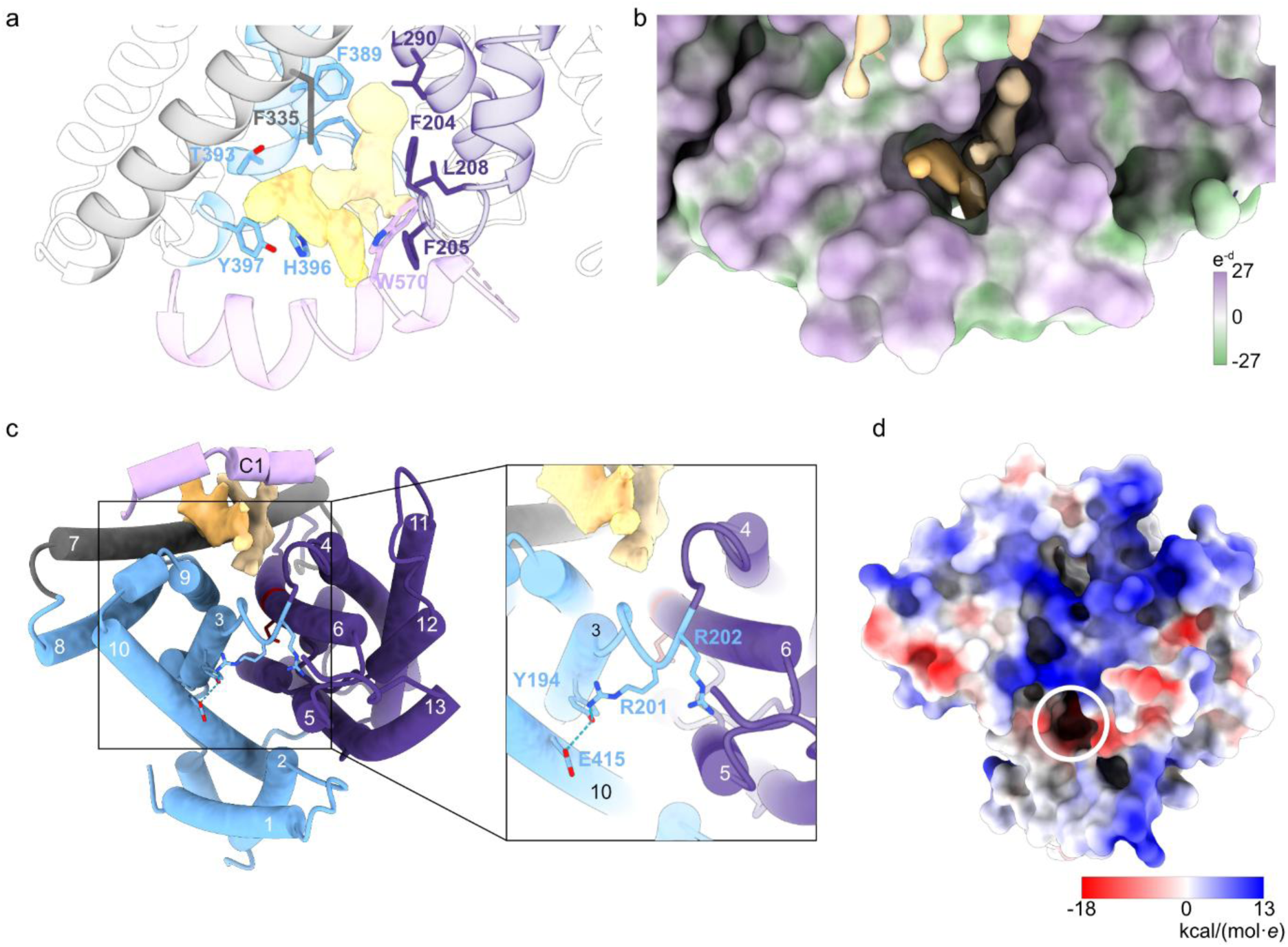
lipids. C-terminus of HsNHE6. **a,** Cartoon representation of the lipid binding cavity with side chain residues forming the cavity and putative lipid density (yellow) **b,** Hydrophobic surface representation of HsNHE6 around the C-terminal helix with putative lipid density (yellow) in the hydrophobic cavity between the helix and the HsNHE6 core. **c,** Cartoon representation of HsNHE6 as seen from the cytosol. The enlarged view shows the positioning of the loop between TM3 and -4 and of R201, R202, Y194 and E415. **d,** Electrostatic surface representation of one HsNHE6 protomer depicted from the cytosolic side. The white circle shows the position of the inward-facing funnel narrowed by the positive charge of R202.

Upon further analysis, the structure of HsNHE6 revealed a distinct hydrophobic cavity on the cytoplasmic side of the protein formed between C1 and TM4, TM6, TM7, and TM9, respectively (**Fig. 3a, b**). Within this cavity, a network of hydrophobic and aromatic residues wraps around cryo-EM densities resembling lipid molecules (**Fig. 3a, b**). To identify the lipid-like densities observed in the cryo-EM map, we performed lipidomics analysis on purified HsNHE6 (**Supplementary Fig. 11**). The most enriched lipid species were phosphatidylcholine (PC) and phosphatidylethanolamine (PE), both of which are abundant components of organelles in the endosomal pathway as well as the plasma membrane^50^. As no lipids were added during purification, the cryo-EM densities and lipidomics data suggest that lipids remained bound to HsNHE6 despite detergent extraction.

Interestingly, the lipid densities are observed to interact with IL2 connecting TM3 and TM4, pushing the loop towards the entrance of the ion-binding site (**Fig. 3c**). The positioning of IL2 is stabilized by hydrogen bond interactions between R201^IL2^ and Y194^TM3^, and between Y194^TM3^ and E415^TM10^. As a result, the positively charged R202^IL2^ is positioned directly at the entrance of the otherwise negatively charged funnel-shaped cavity, thereby narrowing the entrance for ions (**Fig. 3d**). This specific lipid-mediated positioning of a loop has not been observed in other, previously reported NHE structures, which lack defined cryo-EM densities corresponding to lipid molecules at this location and has their IL2 positioned differently than in HsNHE6 (**Supplementary Fig. 12a-d**). Notably, R202^IL2^ is highly conserved among the NHE family members, except for SLC9A8 (**Supplementary Fig. 12 e**), suggesting a potentially conserved regulatory role. R201^IL2^and K200^IL2^ near the entrance to the ion binding site, add to the local positive charge. The positively charged amino acids in these positions are conserved in HsNHE6, -7 and -9.

### The C-terminus of HsNHE6 is intrinsically disordered

As 116 residues composing the distal part of the HsNHE6 C-terminus (G586-A701, termed HsNHE6_G586-A701_ in the following) were unresolved in the cryo-EM structure. Therefore, we experimentally characterized it in isolation by NMR and circular dichroism (CD) spectroscopy. For this, HsNHE6_G586-A701_ was recombinantly expressed and stable isotope-labeled. The ^1^H, ^15^N HSQC NMR spectrum of HsNHE6_G586-A701_ revealed a narrow peak distribution in the ^1^H-dimension, indicative of structural disorder (**Supplementary Fig. 13a**). From the assigned backbone NMR resonances, the extracted secondary chemical shifts (SCS) of C^α^ were all small (SCS for fully formed helix 3.1 ppm). Consecutive positive C^α^ SCSs from T605-N612 and S657-L665 and negative C^α^ SCS values from E614-D620 and L681-D685 reveal lowly populated (<10%) α-helical and extended structures, respectively (**Fig. 4b**). The structural disorder of the HsNHE6_G586-A701_ was further supported by a negative minimum at 205 nm in the far-UV CD spectrum both in the absence and presence of small unilamellar vesicles (SUVs) of different lipid compositions with the intent of providing a membrane surface as a potential interaction partner (**Supplementary Fig. 13b**). These data confirm that the distal part of the HsNHE6 C-terminus is intrinsically disordered in solution, with shorter regions of lowly populated (<10%) helical and extended structures, and without pronounced affinity for the membrane surfaces tested.

**Fig. 4.**
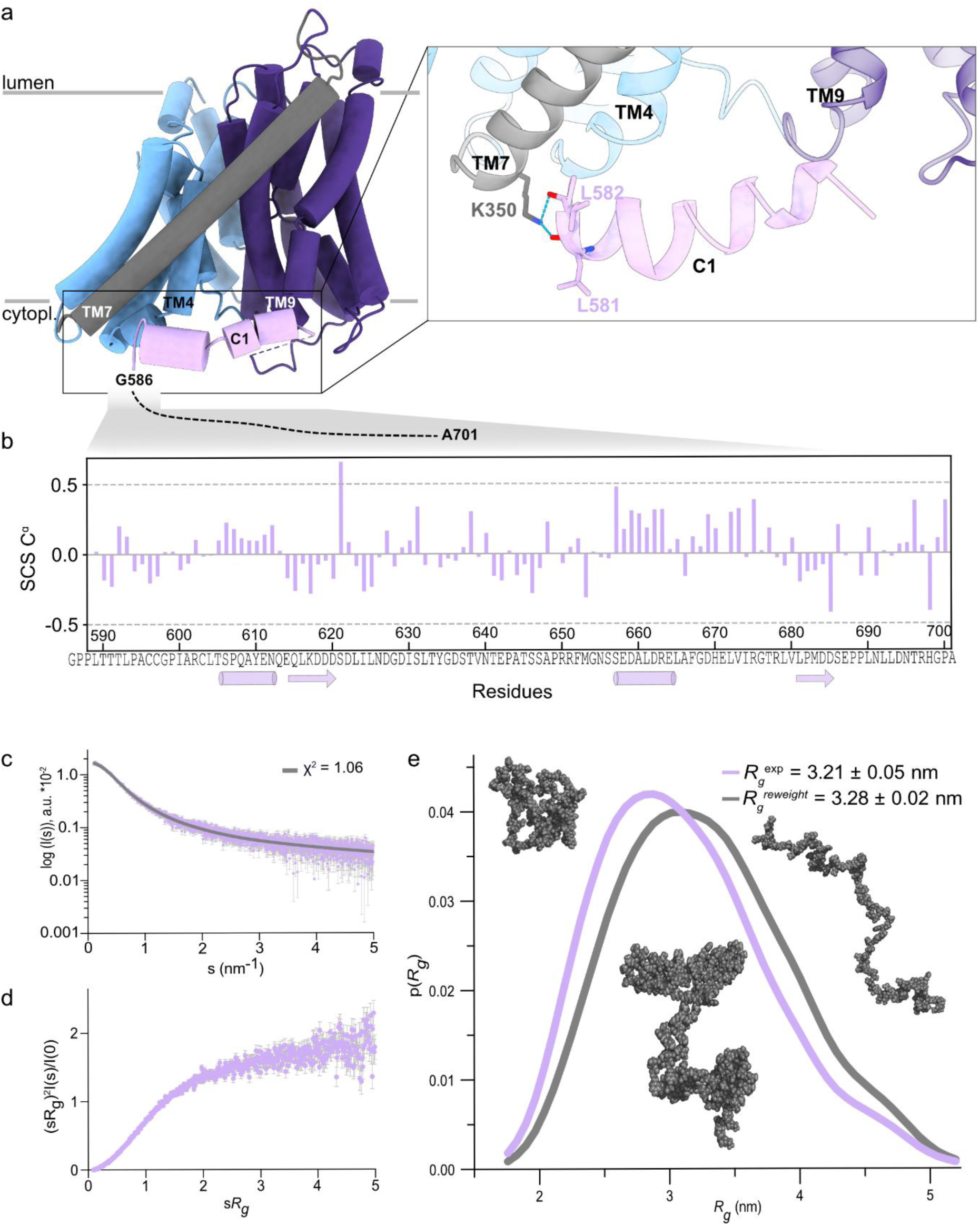
Order and disorder in the C-terminus of HsNHE6. **a,** Cartoon representation of one HsNHE6 monomer with a zoom into the C-terminal C1 helix next to TM4, 7 and 9, with the two hydrogen bonds between K350^TM7^ and the C1 backbone carbonyls. Different domains are colored as in Fig. 1. **b,** NMR secondary chemical shifts (SCS) of C^ɑ^ nuclei for HsNHE6^G586-A701^. The cylinders indicate the positions of transient helices and arrows transient extended structure. **c,** Experimental SAXS scattering data of HsNHE6^G586-A701^ (purple) and back-calculated SAXS scattering curve from the reweighted ensemble from the simulations (grey line). **d,** Dimensionless Krakty plot of the SAXS data. **e,** Probability distribution of *R_g_* of the experimental SAXS data (purple) and from the reweighted ensemble from the simulations (grey). Three structures from the ensemble from the reweighted simulations with different *R_g_*s are represented in a bead-model in grey.

As the C-terminus of HsNHE6 is clearly disordered in solution, we decided to sample the possible configurations of HsNHE6_G586-A701_ using SAXS and coarse-grained molecular dynamics (CG-MD) simulations. The SAXS data on HsNHE6_G586-A701_ shows that it is non-globular according to the upturn in the SAXS Kratky analysis (**Fig. 4c, d**). To obtain an ensemble set of conformations we ran CG-MD simulations of the C-terminal using the CALVADOS model^51^, a one-bead-per-residue model for intrinsically disordered Bayesian maximum entropy. The calculated SAXS intensities showed good agreement with the experimental SAXS data (**Fig. 4c**). From the experimental SAXS data, the *R_g_* was calculated to 3.21 ± 0.05 nm, which is in good agreement with the *R_g_* (3.28 ± 0.02) of the reweighted ensemble. Thus, the SAXS data is consistent with the disordered ensemble for HsNHE6_G586-A701_ (**Fig. 4c**).

### The full-length structural model of HsNHE6 underscores the far-reaching nature of its C-terminus

With the cryo-EM structural model of HsNHE6 and the SAXS supported structure ensemble of the disordered HsNHE6_G586-A701_, it is possible to construct an integrative model of full-length HsNHE6. To represent the breadth of the ensemble of HsNHE6_G586-A701_, we selected three different models distributed across the pair distance distribution range obtained from the SAXS data (**Fig. 4e**). The three representative models of full-length HsNHE6 should not be seen as fully capturing the molecular landscape, and it is possible that the conformations in the crowded cytoplasm may differ from these. However, from the dimensions of the models (up to 130-170 Å maximal IDR expansion) it can be very well appreciated how far the C-terminal region can in principle reach into the cytoplasm and how large the potential for additional protein- and/or lipid interactions of the NHE6 C-terminal region is. It is furthermore likely that some of the more compact states (**Fig. 5**) will make interactions to the core membrane region, and this can potentially interfere with transport both in inhibitory and activating functions.

**Fig. 5.**
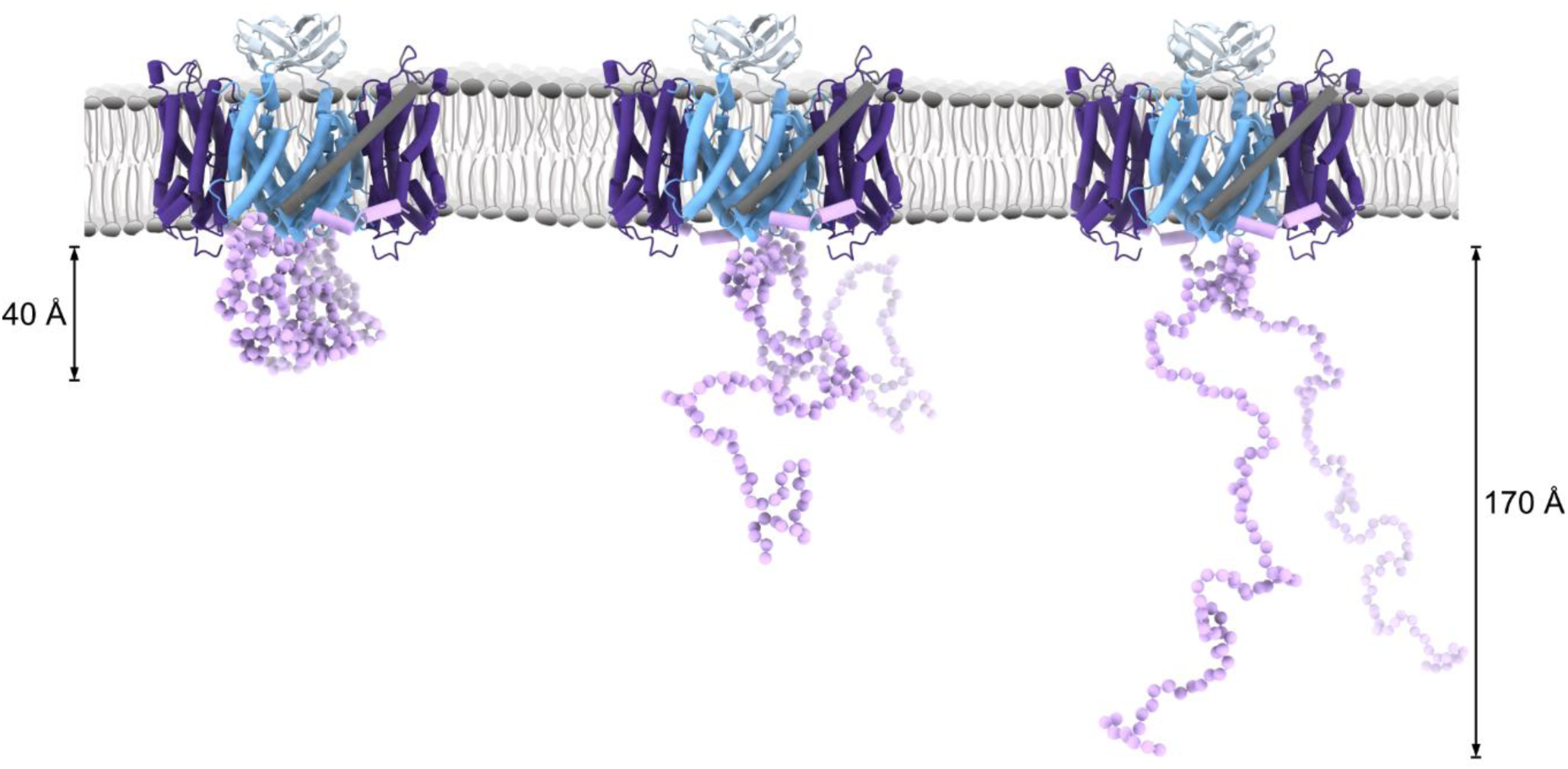
Structural model of full-length HsNHE6. Three different models are shown, representing the structural variability of the C-terminal tail. The models are colored as in Fig. 1, and LL1 between TM2 and - 3 is shown in light blue.

LL1 between TM2 and TM3 of HsNHE6, V122–N170^LL1^, is unique to the endosomal NHEs (NHE6, NHE7, and NHE9). Notably, LL1 is predicted to contain an N-linked glycosylation site at N128^52^, which may contribute to its heterogeneous nature and hinder structural determination. Although of low resolution in the cryo-EM structure, we incorporated a model of LL1 into our integrative model based on a high-confidence AlphaFold3^53^ prediction and the positioning of the homologous loop in EcNHE9^39^ (**Fig. 5**). In both cases, and in accordance with the high content of β-branched amino acid residues as valine and threonine, LL1 is a β-hairpin that interacts with that of the other protomer. Given its proximity to the dimer interface and luminal membrane surface, LL1 is proposed to play a role in protein–protein interactions or lipid modulation in EcNHE9^39^.

### CS mutations locate to sites of ion binding and regulation

Using the structural model of HsNHE6, we mapped CS-linked missense mutations^54,55^ to infer their molecular effects on the function of HsNHE6. As seen from **Supplementary Fig. 14** and **Supplementary Fig. 1**, the pathogenic CS mutations are scattered throughout HsNHE6. While most of the reported mutations are either nonsense or frameshift mutations leading to premature stop codons and non-functional protein^54,56^ (**Supplementary Table 2**), several missense mutations that alter HsNHE6 function have also been reported^54^. One such mutation, G491D^TM12^, is located in the cross over point between TM12 and TM5 and would introduce an additional negative charge at the ion transport site, potentially disrupting ion transport. Another example, L582P^C1^, is located at the interface between TM7 and C1 where a hydrogen bond between the backbone carbonyl of L582^C1^ and K350^TM7^ stabilizes the interaction between TM7 and the C1 helix. The L582P mutation would disrupt this hydrogen bond between TM7 and C1, interfering with the regulatory role of this interaction. In addition, the ΔE287-S288^TM6^ mutations remove two residues close to D292^TM6^ at the ion binding site. Notably, S288^TM6^ is directly involved in ion binding and its loss will therefore very likely interfere with ion engagement and transport, consistent with functional data showing that HsNHE6 carrying this mutation has a reduced capacity for endosomal alkalization^55^.

## Discussion

HsNHE6 is a member of the organellar branch of the NHE family, whose physiological roles and regulatory mechanisms are less understood than those of their plasma membrane counterparts^3,24^. Its critical function in maintaining endosomal pH and trafficking is underscored by its association with CS^32^, a severe X-linked neurodevelopmental and neurodegenerative disorder. However, despite its importance, the structural basis for HsNHE6 function has remained limited.

Our integrative structural and functional analysis provides a comprehensive framework for understanding HsNHE6 and other, organellar Na+(K+)/H+ exchangers. The cryo-EM structure reveals that HsNHE6 adopts the canonical NHE fold of electroneutral NHEs, with 13 TM helices per protomer, forming a homodimeric architecture. The ion-binding site, located at the interface of the core and dimerization domains, is composed of highly conserved residues essential for ion transport. Structural comparisons suggest that HsNHE6 is more similar to the endosomal EcNHE9 than to plasma membrane isoforms such as HsNHE1 or HsNHE3, highlighting potential adaptations to the endosomal-lysosomal environment where organellar NHEs have evolved distinct structural features for regulation specific to the environment.

Our structural and biochemical data revealed that the N-terminus of HsNHE6 undergoes significant co- and post-translational processing. The absence of the first 73 residues in the cryo-EM structure, together with MS identification of L32 as the mature N-terminus, supports the presence of a cleaved signal peptide. This aligns with predictions of ER targeting and suggests that HsNHE6 maturation involves proteolytic processing following N-terminal acetylation. Such modifications may influence protein stability, trafficking, or membrane insertion^57^, and represent a previously uncharacterized regulatory step for HsNHE6 and other NHEs. Following processing at the N-terminus, another key site of modification is found in the LL1 connecting TM2 and TM3, which contains a predicted N-linked glycosylation site^58^ (N128). LL1 was poorly resolved in the cryo-EM structures, likely due to conformational heterogeneity. Interestingly, a shorter splice variant of HsNHE6 lacking most of LL1 (Uniprot ID: Q92581-1; lacking residues 144-175) resembles plasma membrane NHEs and may differ in regulatory interactions, despite similar localization and glycosylation^42,59^. These structural differences raise the possibility of isoform-specific modulation of HsNHE6 function, warranting further investigation.

Cryo-EM analysis revealed distinct densities at the conserved ion-binding site, likely corresponding to bound Na+ and/or water molecules. These densities align with ion-binding positions observed in the crystal structure of PaNhaP^4^ and with MD simulations of EcNHE9^6^, supporting a conserved architecture for monovalent cation coordination. Functional reconstitution of HsNHE6 into BrainPLEx liposomes confirmed its activity as a monovalent cation/H^+^ exchanger, capable of transporting both Na^+^ and K^+^. While previous yeast complementation assays^26^ suggested preference for K^+^, our *in* vitro assays showed no marked selectivity between the two cations. This discrepancy may reflect contextual differences in experimental setup, including membrane composition, curvature, or regulatory context between cellular and reconstituted systems. Together, the structural and functional data underscore the versatility of the HsNHE6 ion-binding site and its capacity to accommodate multiple physiological cations.

Remarkably, a hydrophobic cavity formed by C1 and the TM core raises the possibility for lipid-mediated modulation of HsNHE6. This site includes residues such as D292^TM6^ and S288^TM6^, both critical for ion transport^4,7,8,47^. In addition to structural stabilization, and activation, bound lipids in this site could also contribute to autoinhibition of HsNHE6 or other NHEs. Notably, previous studies proposes that a homologous region to C1 in HsNHE1 can bind lipids such as phosphatidylinositol 4,5-bisphosphate (PI(4,5)P₂), and disruption of lipid interaction significantly reduces HsNHE1 transport activity^60,61^. The presence of a lipid-occupied cavity in HsNHE6 near C1 suggests a similar lipid-based mechanism may exist to restrict or tune activity based on lipid composition, ensuring compartment-specific function. Here, R202^IL2^ may serve as a gating residue that modulates ion access to the binding site. Given the distinct lipid compositions of endosomal versus plasma membranes^62,63^, such as differences in phosphoinositide species or cholesterol content, it is possible that regulation of HsNHE6 activity depends on its subcellular localization, also underscoring the importance of the membrane environment^8,39,61,64,65^.

A distinct feature of HsNHE6 is the presence of C1, a 20-residue C-terminal α-helix, which interacts with TM7 through hydrogen bonding involving the highly conserved K350^TM7^ and the backbone of L581^C1^ and L582^C1^. A similar interaction was recently observed in EcNHE9^39^, suggesting a conserved interaction. So far, such a C-terminal helix has not been described in other mammalian NHEs in the absence of a binding partner. In the case of HsNHE1^7^, it has been proposed that binding of CHP1 to the C-terminal domain modulates activity by favoring an outward-open conformation, potentially counteracting an autoinhibitory effect of the C-terminus^66^. Similarly, it was observed that Ca^2+^-dependent calmodulin (CaM) binding to the C-terminus activates HsNHE1 and releases the autoinhibitory effect^20,21^, underpinning its importance in regulating the functions of NHEs. For HsNHE6, no such binding partners have been confirmed^67^. Thus, it is possible that in HsNHE6 and EcNHE9^39^, the positioning of C1 adjacent to the TM core represents an intrinsic regulatory mechanism, potentially stabilizing the inward-open state and modulating activity in the absence of external regulators. Whether this mechanism is further regulated by specific lipids, post-translational modifications, protonation or other processes, remains to be elucidated.

The distal part of the C-terminus of HsNHE6, unresolved in the cryo-EM structure, is intrinsically disordered and can exist in compact and extended forms, reaching up to 170 Å beyond the endosomal membrane as confirmed by SAXS and MD analyses. By analogy with the multiple interaction partners identified for the C-termini of other NHEs^3^, and like most IDRs^68^, it comprises a platform for interaction partners that may regulate or be regulated by HsNHE6. This region harbors numerous predicted phosphorylation sites and may serve as a scaffold for kinases, phosphatases and other binding partners. Future studies investigating the full interactome of NHE6 will be invaluable for decomposing the regulation of HsNHE6, potentially linking endosomal ion exchange to broader cellular signaling networks.

Importantly, our structural model provides a possibility for interpreting disease-associated HsNHE6 mutations. Several CS-linked missense mutations map to the ion-binding site or to regions involved in structural stabilization, such as the TM7-C1 interface. These mutations likely impair ion coordination or conformational transitions essential for transport. Future mutational and functional studies guided by this model will be critical for elucidating the molecular pathology of CS and identifying therapeutic targets.

In summary, this work presents the full-length integrative structural model of a human NHE, integrating cryo-EM, NMR, SAXS, and functional assays. It establishes a foundation for understanding the mechanistic basis of HsNHE6 function, its regulation by lipids and post-translational modifications, and its disruption in disease. These insights pave the way for future studies into the regulation, interactome, and therapeutic targeting of organellar NHEs.

## Methods

### HsNHE6 construct design

The codon-optimized gene of human NHE6 (HsNHE6, Uniprot ID: Q92581-2; residues 1-701) was purchased from GenScript in a pcDNA3.1(+) vector containing a C-terminal FLAG-tag (HsNHE6-FLAG). For expression in FreeStyle293-F cells, HsNHE6 was cloned into a modified pEG BacMam vector^69^ using restriction-free cloning. The resulting construct contains a C-terminal eGFP-tag followed by a STREPII-tag, separated from HsNHE6 by a TEV protease cleavage site (HsNHE6-GFP-Strep).

### HsNHE6 expression

HsNHE6 was expressed in FreeStyle293-F cells (HEK293F; Gibco) using baculovirus transduction^69^. P1 and P2 viruses were generated in Sf9 insect cells (Gibco). FreeStyle293-F cells were transduced with the P2 virus 24 hours after seeding at a density of 1×10^6^ cells/mL in FreeStyle 293 expression medium (Gibco) supplemented with 2% heat-inactivated fetal bovine serum (FBS, Cytiva). Cells were cultured at 37 °C with 8% CO2 on an orbital shaker. Sodium butyrate was added 8 hours after transduction to a final concentration of 10 mM, and cells were harvested after an additional 40 hours. Cell pellets were snap-frozen in liquid nitrogen for storage until purification.

### HsNHE6 purification in detergent micelles

HsNHE6 was purified at 4 °C. FreeStyle932-F cells expressing HsNHE6 were resuspended in HBS-PI (20 mM HEPES, pH 7.4, 150 mM NaCl, protease inhibitor cocktail) and solubilized with 20 mM DDM/4.1 mM CHS for 1.5 hours under rotation. Insoluble cell debris was removed by centrifugation at 26,000×g for 30 minutes. The supernatant was diluted to 4 mM DDM/0.82 mM CHS with HBS-PI and incubated under rotation with pre-equilibrated StrepTactin Sepharose High 10 Performance resin (Cytiva, Marlborough MA, US) overnight. The beads were washed with 10 bed volumes (BV) of washing buffer 1 (HBS supplemented with 0.085 mM GDN/0.02 mM CHS and 5 mM MgCl_2_, 5 mM ATP), followed by 10 BV of washing buffer 2 (HBS supplemented with 0.085 mM GDN/0.02 mM CHS). The protein was eluted in elution buffer (HBS supplemented with 0.085 mM GDN/0.02 mM GDN-CHS and 5 mM D-desthiobiotin). The eluted protein was subsequently concentrated using a 50-kDa cutoff spin concentrator, and the C-terminal tag was removed by incubation with TEV protease (3:1 molar ratio) overnight. HsNHE6 was further purified by size exclusion chromatography (SEC) using a Superose 6 Increase 10/300 column (Cytiva, Marlborough MA, US) equilibrated in HBS supplemented with 0.034 mM GDN/0.008 mM CHS. Fractions containing the protein were pooled and concentrated. For purification of HsNHE6 in DDM/CHS micelles, GDN/CHS was replaced in all buffers by 0.5 mM DDM/0.1 mM CHS.

### Cryo-EM sample preparation and data acquisition

HsNHE6 structure in GDN/CHS detergent micelles. Purified HsNHE6 (3 μl of 2.63 mg/ml) was applied to freshly glow-discharged Quantifoil Holey Carbon Films R 1.2/1.3 on AU 300 mesh grids. The grids were blotted for 5.5 s at 4 °C under 100% humidity using a Vitrobot Mark IV (Thermo Fisher Scientific^TM^) and flash-frozen in vitreous liquid ethane. Cryo-EM data were collected on a Titan Krios G2 Cryo-TEM equipped with a Thermo Scientific^TM^ Selectris X imaging filter and a TFS Falcon IVi detector operated at 300 keV. The videos were collected at a nominal magnification of 165,000, corresponding to a pixel size of 0.725 Å. Data were collected with a defocus range of 0.8 to 2.5 µm and a total does of 49 e^-^/A^2^ and images were saved as Electron Event Recordings (EER).

### Cryo-EM image processing

Image processing of HsNHE6 in GDN/CHS micelles was carried out in RELION-5.0-beta^70^. Movies were motion corrected using Relion’s own implementation of MotionCor2^71^, the contrast transfer function (CTF) was estimated by CTFFIND-4.1^72^ and image processing was continued with movies with an estimated CTF below 5 Å. From 4.507 micrographs, about 2,000 particles were extracted after manual picking and underwent two rounds of 2D classifications. Particles from good classes were used to train Topaz^73^ and 732,718 particles were extracted after Topaz picking from all micrographs. These particles underwent another 4 rounds of 2D classification and particles from good classes were used for *ab initio* reconstruction to generate an initial model. Subsequently, particles underwent 3D classification with Blush regularization^74^ into 4 classes without imposing symmetry and the map form the best class was refined with Blush regularization either without imposing symmetry or with imposed C2 symmetry. After another round of 3D classification without image alignment, particles from the best respective class were refined in 3D using Blush regularization followed by post-processing with a mask resulting in reconstructions of resolutions 3.5 Å (C1) and 3.4 Å (C2), respectively. Details regarding image processing are shown in **Supplementary Fig. 2, 3, 5** and **Supplementary Table 1.**

### Model building and refinement

*HsNHE6 structure in GDN/CHS detergent micelles.* Model building into the C2 cryo-EM map was carried out using the automated model building neuronal network ModelAngelo^41^. Residues missing from automated model building were built using Coot^75^. The model was refined and validated using Phenix and MolProbity^76,77^. For the C1 cryo-EM map, Namdinator^78^ was used to fit the refined C2 model, followed by manual examination in Coot^75^ and further refinement and validation using Phenix and MolProbity^76,77^. An overview of the residues included in the final refined models including their refinement statistics and validation are shown in **Supplementary Table 1** and **Supplementary Fig. 1**. Figures are prepared with UCSF ChimeraX^79^.

*HsNHE6 full-length model.* To generate a full-length model of HsNHE6, the unresolved loop between TM2 and -3 was modeled using the highest-confidence AlphaFold3^40^ prediction. This loop was manually integrated into the cryo-EM structural model and the connecting peptide bond between the structural model and the grafted loop segment was refined using Isolde^80^. For the C-terminus, three representative models were selected from the HsNHE6 ensemble generated from SAXS data, reflecting a range of conformational states of the tail, from fully compact to fully extended. Each conformation was individually grafted onto the HsNHE6 structural model, and the peptide bonds between the core model and the appended tails were refined using Isolde^80^.

### Expression and purification of *E. coli* F1F0-ATP synthase

*E. coli* F1F0-ATP synthase was expressed and purified as described previously^81^. After expression in *E. coli* BL21 (DE3) pLysE cells, cells were disrupted by sonication and membranes were harvested. The protein was extracted from the membranes with extraction buffer (50 mM Tris/HCl, pH 7.5, 100 mM KCl, 250 mM sucrose, 40 mM ε-aminocaproic acid, 15 mM p-amino-benzamidine, 5 mM MgCl_2_, 0.1 mM EDTA, 0.2 mM DTT, 0.8% phosphatidylcholine, 1.5% octyl glucoside, 0.5% sodium deoxycholate, 0.5% sodium cholate, 2.5% glycerol, and 30 mM imidazole) by incubating for 1.5 h on a tube roller and cell debris was afterwards removed by ultracentrifugation. The supernatant was loaded on a 1 ml HisTrap FF column (Cytiva, Marlborough MA, US), and the column was washed with 15 column volumes (CV) extraction buffer, before elution of the protein in 3 ml elution buffer (extraction buffer supplemented with 180 mM imidazole).

### Coupled-proton transport assay

Brain Polar Lipid Extract (BrainPLEx, Avanti Research) solubilized in chloroform were dried under a constant stream of N_2_ and CHS powder was added in a 5:1 w/w ratio. Lipids and CHS were resuspended in 30 mM HEPES (pH 8.0), 150 mM RbCl, 5 mM MgCl_2_ to a final lipid concentration of 10 mg/ml. The lipid solution underwent four rounds of sonication in a bath sonicator before the liposomes were subjected to extrusion using a polycarbonate filter with a pore size of 100 nm. The reconstitution of protein into the BrainPLEx/CHS liposomes was performed based on the previous published protocol^45^. Briefly, liposomes were destabilized with Na-cholate before the addition of 100 µg of HsNHE6 and *E. coli* F0F1 ATP synthetase (2:1 molar ratio). After incubation for 30 min at room temperature, detergent was removed using PD-10 desalting columns. For the functional assay, 10 µl of protein containing liposomes were diluted into 100 µl of assay buffer (30 mM HEPES (pH 8.0), 150 mM RbCl, 5 mM MgCl_2_, 5 µM ACMA and 130 nM valinomycin) in a 96-well plate. Upon the addition of 2.5 mM ATP, an outward-directed pH gradient was built-up, and the transport was initiated by the addition of NaCl after equilibration (∼3 min). After 60 s, the proton gradient was dissipated by the addition of 20 mM NH_4_Cl. The experiments were carried out in a FlexStation 3 Multi-Mode Microplate Reader (Molecular Devices, San Jose CA, US) and the pH-dependent quenching and dequenching of ACMA fluorescence at 480 nm (excitation wavelength: 410 nm) was recorded. To assess the cation dependence of HsNHE6, the experiments were also conducted upon the addition of KCl instead of NaCl. The fluorescence data were normalized, and data are presented as the mean of three independent purifications, with the standard deviation (SD) representing the variability within the samples.

### Expression and purification of HsNHE6_G586-A701_

The distal part of the NHE6 C-terminus, HsNHE6_G586-A701_, was produced with an N-terminal hexa-histidine small ubiquitin-like modifier (H_6_-SUMO) tag, which can be cleaved off with ubiquitin-like protein protease 1 (ULP1). The coding region was inserted into a modified pET24a vector (Twist Bioscience, US). The protein was expressed in *Escherichia coli* (*E. coli*) BL21(DE3) cells and cultured in Luria Bertani (LB) broth medium, or M9 minimal medium containing ^15^N-NH_4_Cl and ^13^C_6_-glucose, at 37°C. Protein expression was induced at OD_600_ 0.6-0.8 with 0.5 mM isopropyl-*β*-D1-thiogalactopuranoside (IPTG) and the cells grown for 4 hours at 37°C before being harvested by centrifugation (5,000 x g, 15 min, 4°C). The cell pellet was resuspended in Buffer A (50 mM Tris-HCl, pH 8.0, 150 mM NaCl, 10 mM imidazole, 1 mM DTT), and lysed by French Press at 25 kPsi on a Constant System Ltd MC cell Disruptor (Contant Systems, Daventry, UK). The soluble fraction was collected by centrifugation (20,000 x g at 4°C for 30 min) and applied on a 3 mL pre-equilibrated Nickel Sepharose Fast Flow resin (Cytiva, Marlborough MA, US). The column was washed with 10 x column volume (CV) of Buffer B (50 mM Tris-HCl, pH 8.0, 1 M NaCl, 10 mM imidazole, 1 mM DTT), and 10 X CV Buffer A, before the protein was eluted with 15 mL Buffer C (50 mM Tris-HCl, pH 8.0, 150 mM NaCl, 250 mM imidazole, 1 mM DTT). ULP1 was added to the eluted fraction, the solution was dialyzed against Buffer A overnight at 4°C and subsequently loaded onto a 3 mL pre-equilibrated Nickel Sepharose Fast Flow resin (Cytiva, Marlborough MA, US). The flowthrough was either concentrated to 2 mL using a 3-kDa cutoff Amicon® centrifugal filter device (Merck, Darmstadt, Germany) or directly applied to a pre-equilibrated (50 mM NH_4_HCO_3_ pH 7.8) 3 mL Source 15RPC column (Cytiva, Marlborough MA, US). The protein was eluted using a 0-100 % (v/v) gradient of 50 mM NH_4_HCO_3_ pH 7.8, 70 % (v/v) acetonitrile. Lastly, the protein was lyophilized to allow solubilization in experiment-appropriate buffer.

### CD spectropolarimetry of HsNHE6_G586-A701_

Far-UV circular dichroism (CD) spectra ware recorded on Jasco J-815 CD spectropolarimeter using a 1 nm bandwidth, a digital integration time (D.I.T.) of 1-4 seconds), a scan speed of 50 nm/min, and 10 accumulations. Background spectra recorded identically were subtracted and only measurements with a high-tension (HT) voltage below 700 V were shown. Lyophilized HsNHE6_G586-A701_ was resolubilized in 10 mM NaH_2_PO_4_/Na_2_HPO_4_ pH 7.3, 150 mM NaF, alone or with the inclusion of 0.5 mM POPC SUVs or 0.5 mM POPC:POPS (3:1) SUVs (Avanti Polar Lipids). The lipids were dissolved in chloroform, and for POPC to POPS mixed in a 3:1 ratio before evaporating the organic solvent with a stream of N_2_ and resolubilizing the lipids for extrusion.

### NMR Spectroscopy of HsNHE6_G586-A701_

Double-labelled (^15^N,^13^C) lyophilized HsNHE6_G586-A701_ was resolubilized in 20 mM NaH_2_PO_4_/Na_2_HPO_4_, pH 7.4, 150 mM NaCl, 5mM DTT, 10 % (v/v) D_2_O, 250 µM 4,4-dimethyl-4-silapen-tane-1-sulfonic acid (DSS) in volume of 600 µL. Before transferred to a single use 5 mm Bruker Labscape Essence 7” NMR tube, the sample was pH adjusted and centrifuged for 10 min at 4°C and 20,000 x g. Spectra for backbone assignment of HsNHE6_G586-A701_ (500 µM) were acquired on a Bruker Avance III HD 750 MHz spectrometer equipped with a 5 mm TCI cryoprobe. The assignments were done manually based on the analysis of 2D ^1^H, ^15^N-HSQC, and 3D HNCACB, HN(CO)CACB, HNCO, HN(CA)CO, and HN(CA)NNH experiments obtained with non-uniform sampling and recorded at 5 °C. Raw free induction decays were processed and visualized using NMRpipe^82^ or Topspin 3.7.0^83^ and analyzed in CcpNMR Analysis 2.5.2^84^. Chemical shifts were referenced to the DSS signal^85^ and indirectly using gyromagnetic ratios. The secondary chemical shift (SCS) of C^α^ for each residue was calculated by subtracting the observed δC^α^ from of the sequence corrected random-coil chemical shift (POTENCI^86^) value.

### SAXS of HsNHE6_G586-A701_

Lyophilized unlabeled HsNHE6_G586-A701_ was dissolved in 20 mM HEPES, pH 7.4, 150 mM NaCl, 5 mM DTT, and further dialyzed overnight at 4°C against the same buffer. A concentration series of the protein was made in a range from 1 to 4 mg/mL. Small angle X-ray scattering was performed at the EMBL P12-bioSAXS beamline at PETRAIII (DESY, Hamburg, Germany)^87^. Data was collected on a Pilatus 6M detector with a distance of 3 meters. Samples were loaded automatically using a sample changer. Scattering intensities were measured as a function of the momentum transfer s = (4πsin*θ*)/λ where 2*θ* is the scattering angle and λ is the X-ray wavelength (0.12398 nm) at 20 °C. The exposure time was 0.095 s, and the measured angles were from 0.000225 to 0.0728 nm. Buffer measurements were subtracted to get the final data files. Primary data analysis and Guinier analysis were done using PRIMUS^88^. For the final analysis only the data from the concentration of 4 mg/mL HsNHE6_G586-A701_ was used.

### Molecular dynamics simulations of HsNHE6_G586-A701_

To obtain an ensemble of the HsNHE6_G586-A701_ protein, CALVADOS 2^51^ was used. The input parameters were the amino acid sequence, pH 7.4, 0.15 M NaCl, and 20 °C, as well as the SAXS scattering curve.

### IFM analysis of HEK293 cells expressing HsNHE6

FreeStyle293-F cells were seeded 24 h pre-transfection in Freestyle medium supplemented with 2% FBS onto poly-L-lysine coated cover slips in 6-well plates (2 ml of cells per well, 0.5×10^6^ cells/ml). On the day of transfection, 1 μg of DNA (HsNHE6-FLAG in pcDNA3.1(+) vector) was incubated with 1 μl of FectoPro in medium without FBS for 2 min and then added to the cells. 48 h after transfection, cells were washed with ice-cold PBS, fixed with 4% Paraformaldehyde (VWR) and kept in TBS-T. After fixation, cells were permeabilized in 0.5 % Triton-X-100 (X100, Sigma Aldrich), blocked in 5% BSA (A7906, Sigma Aldrich) in 1X TBST, and incubated with primary diluted in 1% BSA in 1X TBST for 1.5 h at room temperature. Thereafter, coverslips were washed thrice in 1X TBST and incubated with secondary antibodies (1:600), and if indicated, with rhodamine phalloidin (1:1000), diluted in 1% BSA (A7906, Sigma Aldrich) in 1X TBST for 1.5 h at room temperature. Coverslips were counterstained with DAPI (1:1000), washed thrice with 1% BSA (A7906, Sigma Aldrich) in 1X TBST and mounted in N-propyl-gallate mounting medium. Cells were imaged at 100 X magnification using an Olympus IX83 spinning-disk confocal microscope with a Yokogawa scanning unit. Images were processed using ImageJ software. Antibodies used for IFM analysis are listed in **Supplementary Table 3**.

### Lipidomics

Lipidomics analyses were performed on purified HsNHE6 (6.5 µM) or DDM/CHS-solubilized cells either overexpressing HsNHE6 or not (125 mg/ml cells) as described previously^89^. Briefly, samples were extracted using Folch extraction^90^. Prior to tissue lysis, Splash mix (Merck) was added to the extraction solvent. After centrifugation and phase separation, the apolar and polar phases were transferred to separate tubes, and the apolar phase dried under N_2_. The apolar phase was resuspended in 30 µl methanol/chloroform (1:1) and centrifuged (5 min, 16,000 × *g*, 22 °C) before transferring to HPLC vials. A quality control sample was constructed by pooling 3 µl of each sample. Samples (0.5 µl) were injected using a Vanquish Horizon UPLC (Thermo Fisher Scientific) equipped with a Waters ACQUITY Premier CSH (2.1 × 100 mm, 1.7 µM) column operated at 55 °C. The analytes were eluted using a flow rate of 400 μL/min and the following composition of eluent A (Acetonitrile/water (60:40), 10 mM ammonium formate, 0.1% formic acid) and eluent B (Isopropanol/acetonitrile (90:10), 10 mM ammonium formate, 0.1% formic acid): 40% B from 0 to 0.5 min, 40–43% B from 0.5 to 0.7 min, 43–65% B from 0.7 to 0.8 min, 65–70% B from 0.8 to 2.3 min, 70–99% B from 2.3 to 6 min, 99% B from 6–6.8 min, 99–40% B from 6.8–7 min before equilibration for 3 min with the initial conditions. The flow from the UPLC was coupled to a TimsTOF Flex (Bruker) instrument for mass spectrometric analysis. Compounds were annotated in Metaboscape (Bruker) using both an in-built rule-based annotation approach and using the LipidBlast MS2 library^91^. Features were removed if their average signal were not >5× more abundant in the QC samples than blanks (water extraction).

### LC-MS/MS analysis of HsNHE6

For LC-MS/MS analysis, 25-30 μg of purified HsNHE6 in 20 mM HEPES (pH 7.4), 150 mM NaCl, 0.5 mM DDM/CHS were prepared using S-trap and trypsin digestion. Trypsin cleaves peptide bonds at the carboxyl side of lysine and arginine residues, except when followed by proline^92^. The sample was analyzed using an Orbitrap Eclipse mass spectrometer with an Easy 1200 nUPLC system and data were searched against an in-house database containing the target protein using the Mascot search engine. Semitryptic peptides, peptides with one terminus consistent with trypsin cleavage and the other resulting from non-tryptic processing, were also considered in the search to account for potential endogenous or partial proteolytic events e.g. due to lack of accessibility.

## Supplementary information

The accompanying supplementary figures and tables are provided as a separate file.

## Acknowledgements

The authors thank Casper de Lichtenberg for assistance with freezing cryo-EM grids, and Tillmann H. Pape and Nicholas Heelund Sofos from the Core Facility for Integrated Microscopy (CFIM, University of Copenhagen) for their support during data collection. We are also grateful to Signe A. Sjørup for technical support in preparing the C-terminus, and to beamline scientist Melissa Ann Graewert at EMBL Hamburg for her assistance with SAXS measurements at PETRA III, conducted under the BAG proposal SAXS-1342. Technical support on the IFM was kindly provided by Mette Flink. Expression of HsNHE6 was carried out using a modified BacMam vector, generously provided by Eric Gouaux (Addgene plasmid #160680). The pFV2-HA plasmid was a gift from Steven Vik (Addgene plasmid #187082; http://n2t.net/addgene:187082; RRID: Addgene_187082). The authors thank Ciara F. Pugh and Maria F. Vicino for fruitful discussions and input for this manuscript.

## Declarations

### Inclusion and ethics

In preparing this manuscript, authors ensured equitable recognition of all contributors. This study did not involve human participants, human tissues, or animal research requiring ethical approval. The research adheres to Nature Portfolio’s policies on integrity, accessibility, and reproducibility, ensuring transparency and rigor in scientific reporting.

### Data availability

The reconstructed maps are available from the EMDB under access code EMD-54418 (HSNHE6 in C1 symmetry) and EMD-54419 (HSNHE6 in C2 symmetry). The atomic models are available from the PDB under access codes PDB 9S0R (HsNHE6 in C1 symmetry) and 9S0S (HsNHE6 in C2 symmetry). The raw cryo-EM movies are available under the access code EMPIAR-#####. Chemical shifts of HsNHE6^G586-A701^ have been deposited in the BioMagResBank under the accession code ####. SAXS data are deposited in SASDB under the accession code 6943. Source Data are provided as a Source Data file.

### Contributions

All authors made significant contributions to this research, fulfilling the criteria for authorship. HEA, BBK and SFP conceived the project. HEA built the experimental setup for structural and functional characterization of HsNHE6. LPF carried out HsNHE6 and ATPase expression and purification, the liposomal activity assay, single particle analysis, and model building. JOS expressed and purified the ATPase for the liposome activity assay. JFH and NJKF carried out lipidomics. JFH, NJKF, and LPF processed and analyzed lipidomics data. EET and BBK designed C-terminal HsNHE6 constructs. EET and MRL expressed and purified HsNHE6_G586-A701_ for CD spectropolarimetry, NMR and SAXS. EET, MRL and BBK carried out NMR analysis of HsNHE6_G586-A701_. MRL performed SAXS and MD simulations of HsNHE6_G586-A701_ and analyzed the data with BBK. LPF seeded cells for IFM. LS carried out IFM and analyzed the data with SFP. LPF, LS and MRL prepared figures. LPF and HEA wrote the initial draft of the manuscript. All authors reviewed, provided feedback on, and approved the final manuscript.

### Corresponding authors

Correspondence to Henriette E. Autzen

## Supplementary Information

**Supplementary Fig. 1.**
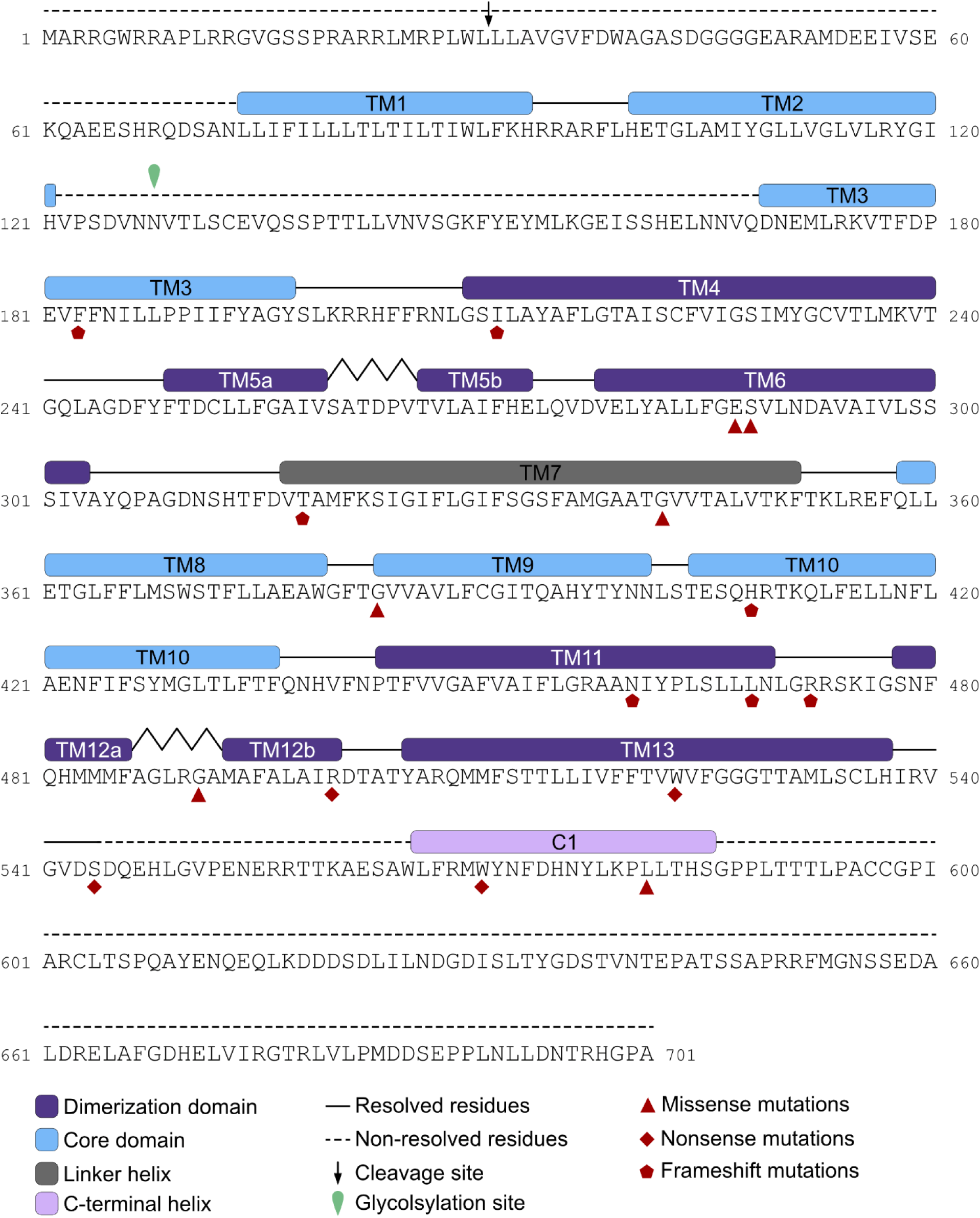
Primary structure of HsNHE6. Dimerization domain TMs (blue), core domain TMs (violet), linker helix TM7 (grey), C-terminal α-helix C1 (pink) are indicated. Resolved and model loop residues are shown as lines, non-modelled residues in dashed lines. Christianson syndrome (CS)-causing mutations reported as either missense (triangle), nonsense (diamond) and frameshift (polygon) are marked, respectively. The arrow indicates the signal peptide cleavage site found through LS-MS/MS analysis in this study, while the green drop marks the reported glycosylation site^1^.

**Supplementary Fig. 2.**
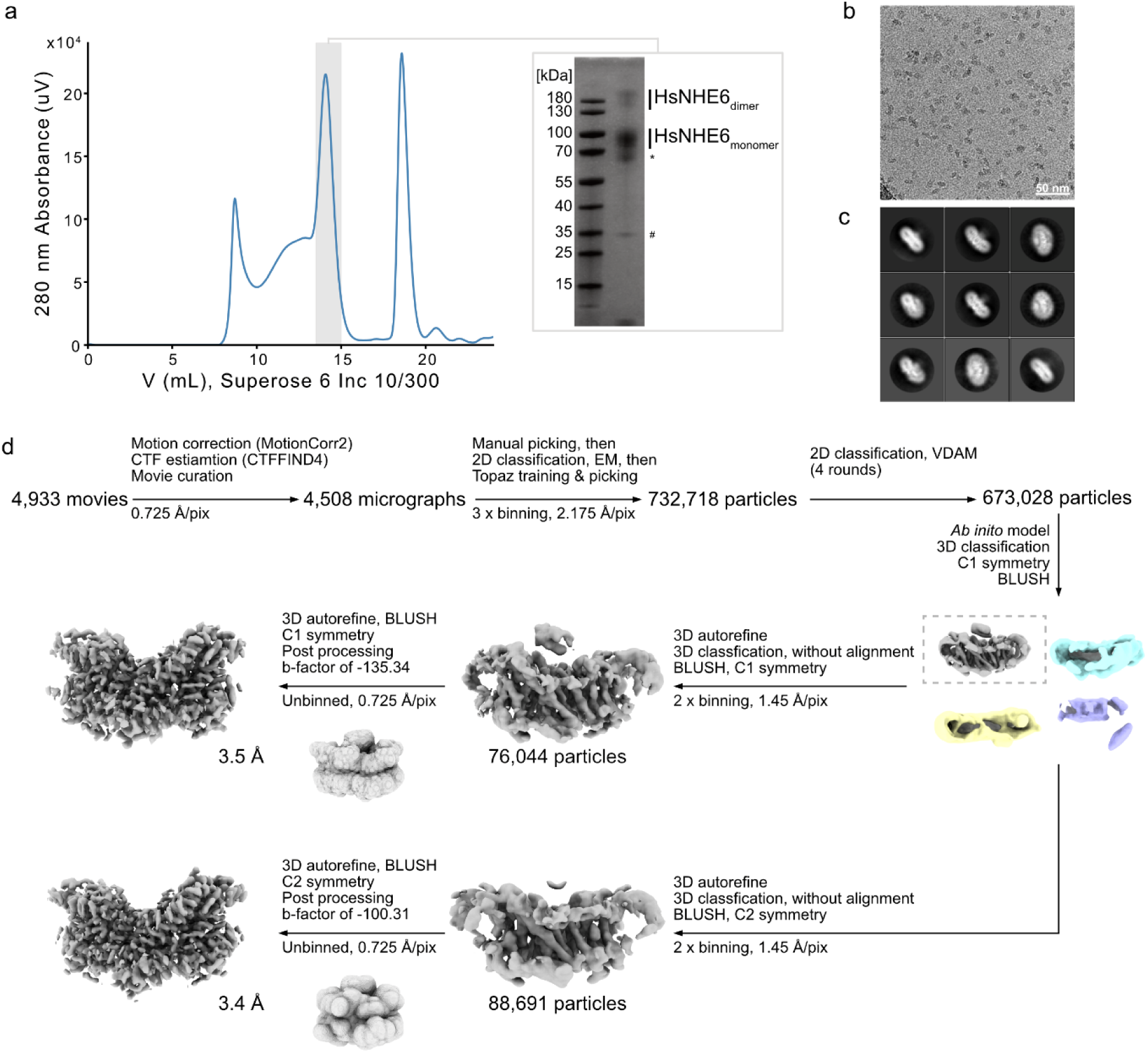
Purification and cryo-EM processing of HsNHE6. **a,** Size-exclusion chromatogram (Superose 6 Increase 10/300) of HsNHE6 in detergent micelles. The fractions collected for cryo-EM sample preparation are indicated by the gray shaded area. An SDS-PAGE gel stained with Coomassie shows the concentrated sample from SEC used for cryo-EM; the asterisk marks a co-purifying heat shock protein, and the pound sign co-purified, cleaved GFP-Strep-tag. **b,** Representative motion-corrected micrograph imaged at 165,000x magnification, the scale bar corresponds to 50 nm. **c,** Representative 2D class averages. **d,** Cryo-EM data processing pipeline for HsNHE6 in GDN/CHS detergent micelles. Processing was carried out in RELION-5.0-beta^2^ and Blush regularization^3^ was applied during 3D classification and refinement steps.

**Supplementary Fig. 3.**
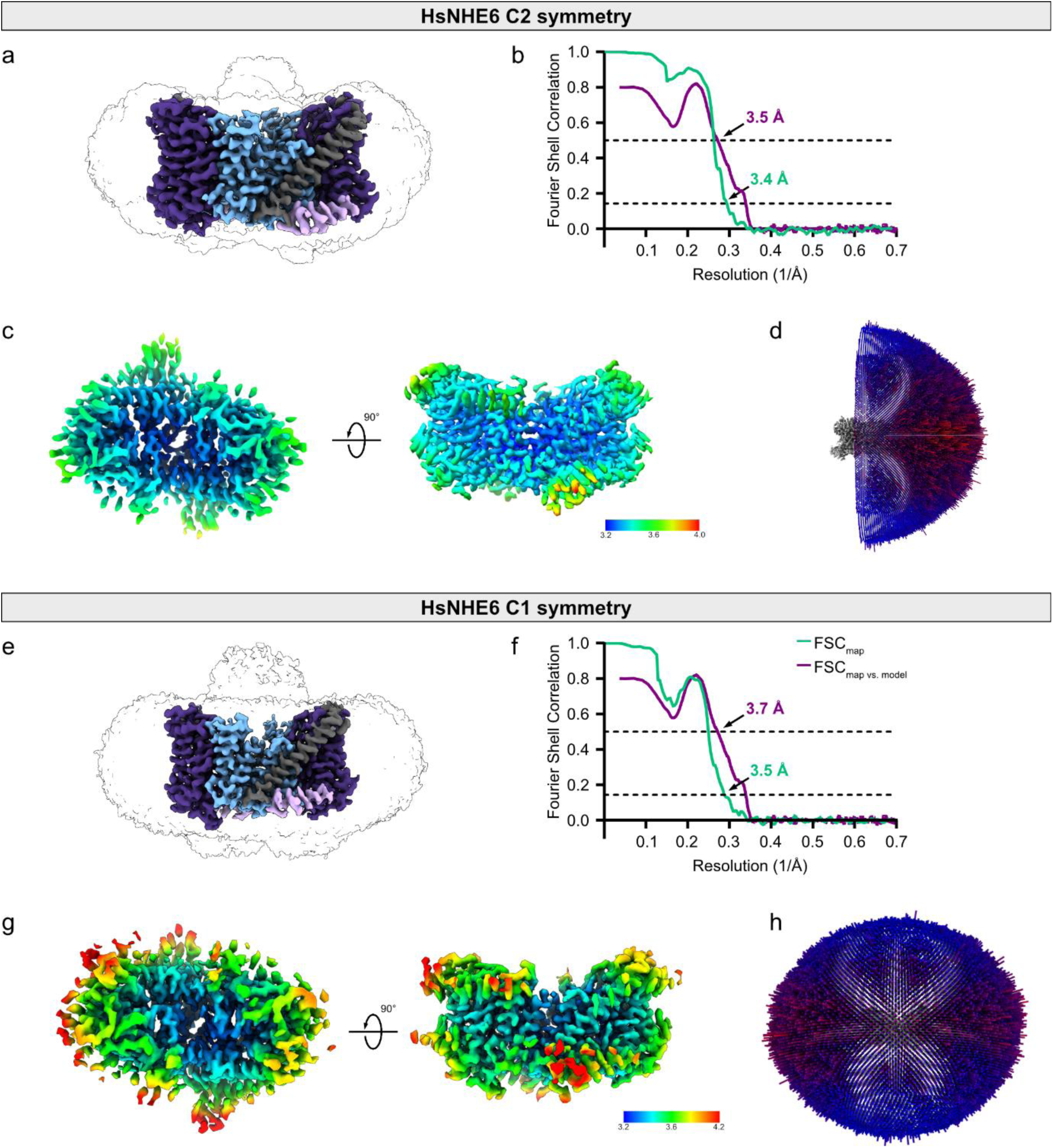
Map and model statistics. **a, e,** Side view of sharpened and unsharpened cryo-EM density maps of HsNHE6 in GDN/CHS micelles. The sharpened map is colored according to the domain organization of the protomers as in previous figures; dimerization domain (blue), core domain (violet), TM7 (gray), C1 helix (pink). The unsharpened map is transparent with a black outline. **b, f,** Fourier shell correlation (FSC) half-map (green) and model-map curves (violet), adopted RELION-5.0-beta^2^ and Phenix Mtriage^4^, respectively. The resolution at FSC = 0.143 for the map and the FSC = 0.5 for the map vs. model are indicated by arrows, respectively. **c, g,** Local resolution representation of the best reconstruction. **d, h,** Angular distribution of particles used in the final reconstruction.

**Supplementary Fig. 4.**
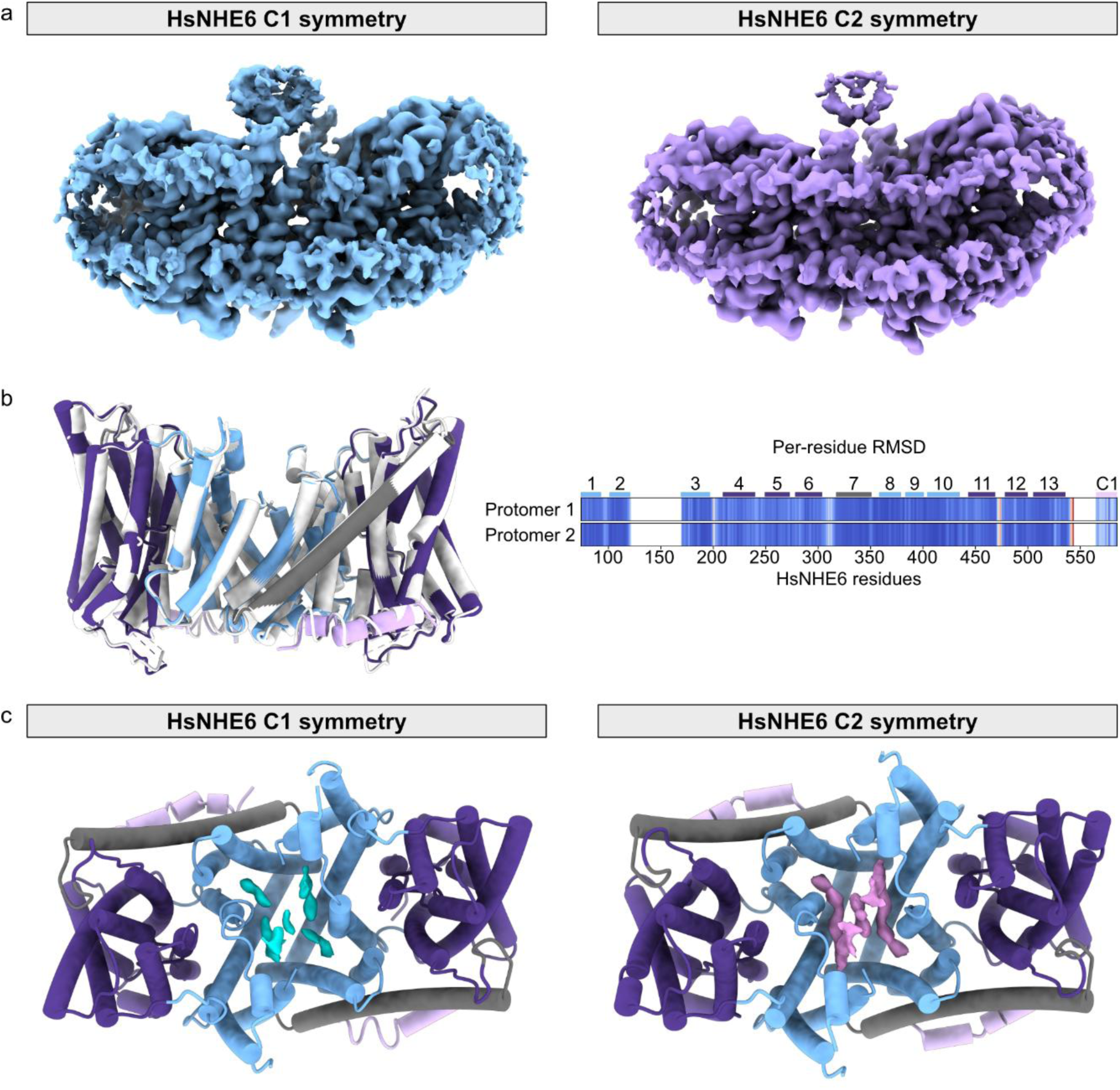
Comparison of structures in C1 and C2 symmetry. **a,** Side view of HsNHE6 structures solved with C1 symmetry (left) and C2 symmetry (right), **b,** Overlay of the HsNHE6 structural models built into the C2 (colored) and C1 (white) maps, respectively. The per-residue RMSD plots show the difference between the Cα atoms for each protomer, after alignment of the core domain TMs. White areas represent regions that are missing in the models, **c,** Lipid densities at the dimer interface of HsNHE6 resolved in the cryo-EM structure in C1 (cyan, left) and C2 symmetry (pink, right). The dimer domain (TM1-3 and TM8-10) is colored light blue, the core domain (TM4-6 and TM11-13) is colored dark purple, the linker helix (TM7) is colored gray, and C1 is colored light purple.

**Supplementary Fig. 5.**
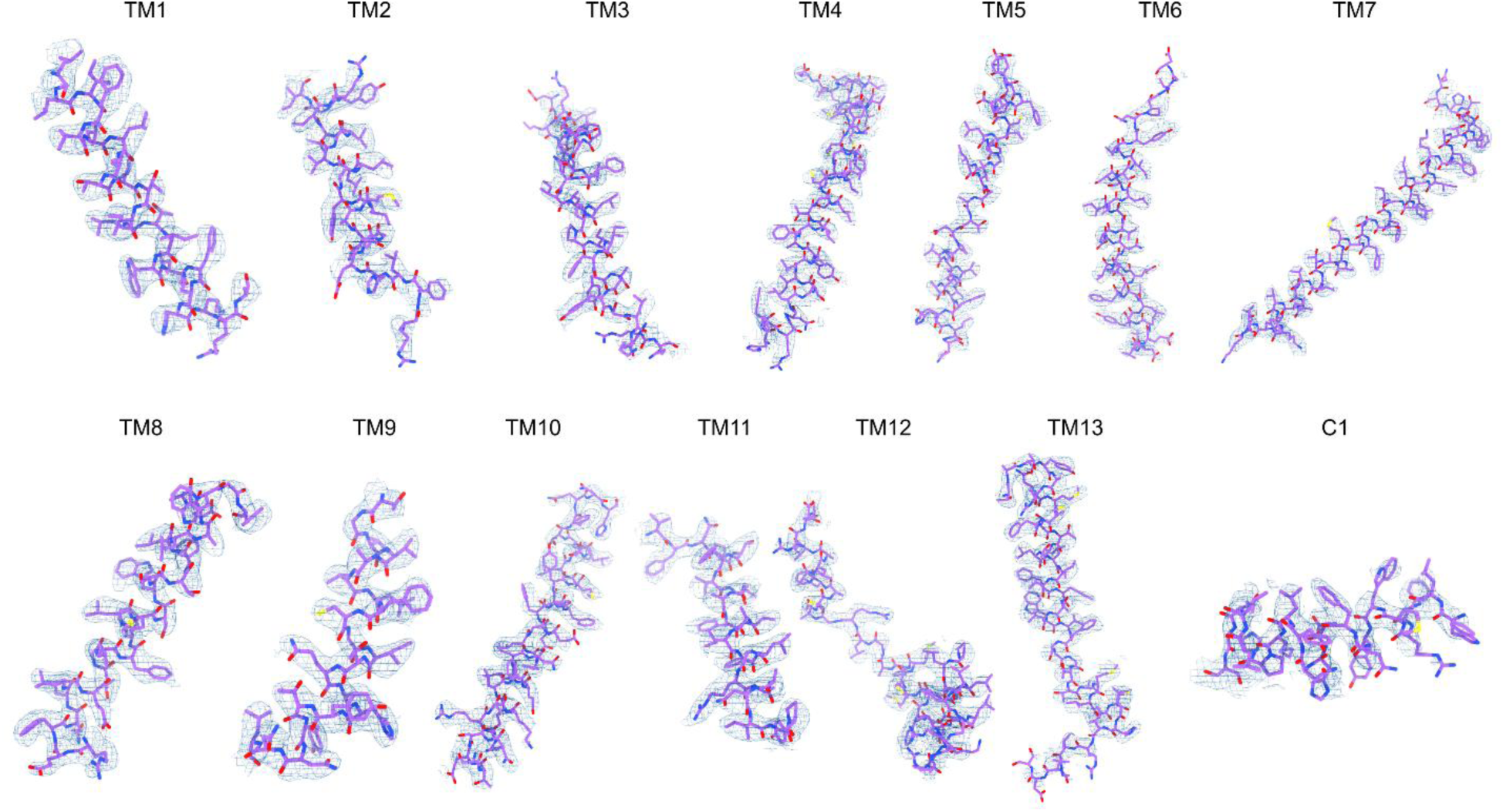
Representative regions of the cryo-EM structure of HsNHE6 in detergent. Representative segments of the HsNHE6 structural model built into the cryo-EM map.

**Supplementary Fig. 6.**
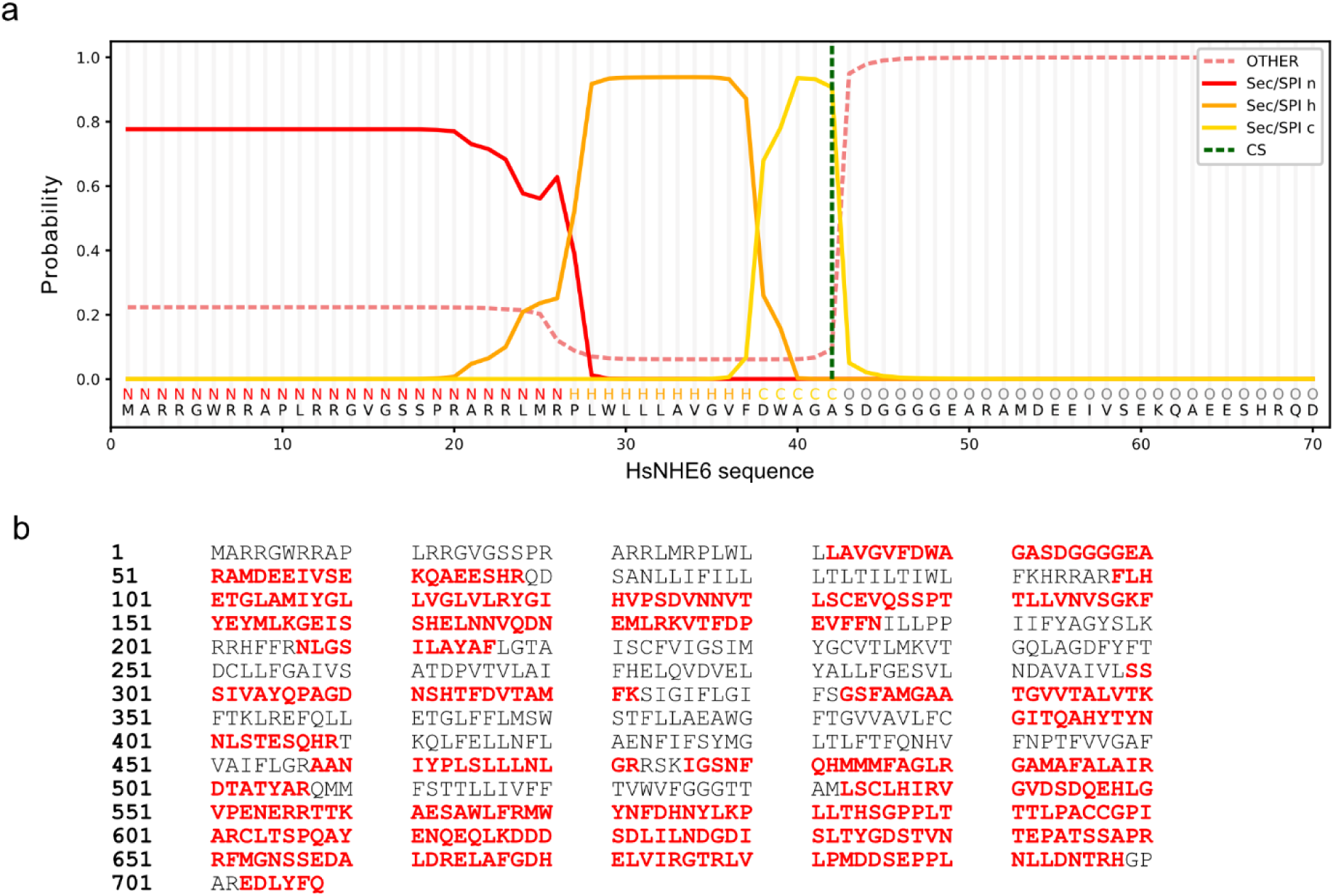
N-terminus of HsNHE6. **a,** Signal peptide prediction for HsNHE6 using SignalP 6.0^5^, indicating a signal peptide between P27 and F37. **b,** Matched peptides (bold red) found after tryptic digestion and LC-MS/MS analysis of HsNHE6 mapped on protein sequence. Semitryptic peptides, defined by one tryptic and one non-tryptic terminus, were included to capture potential endogenous or partial cleavage events due to limited protease accessibility. The most N-terminal peptide is the semi tryptic peptide 32-LAVGVFDWAGASDGGGGEAR-51.

**Supplementary Fig. 7.**
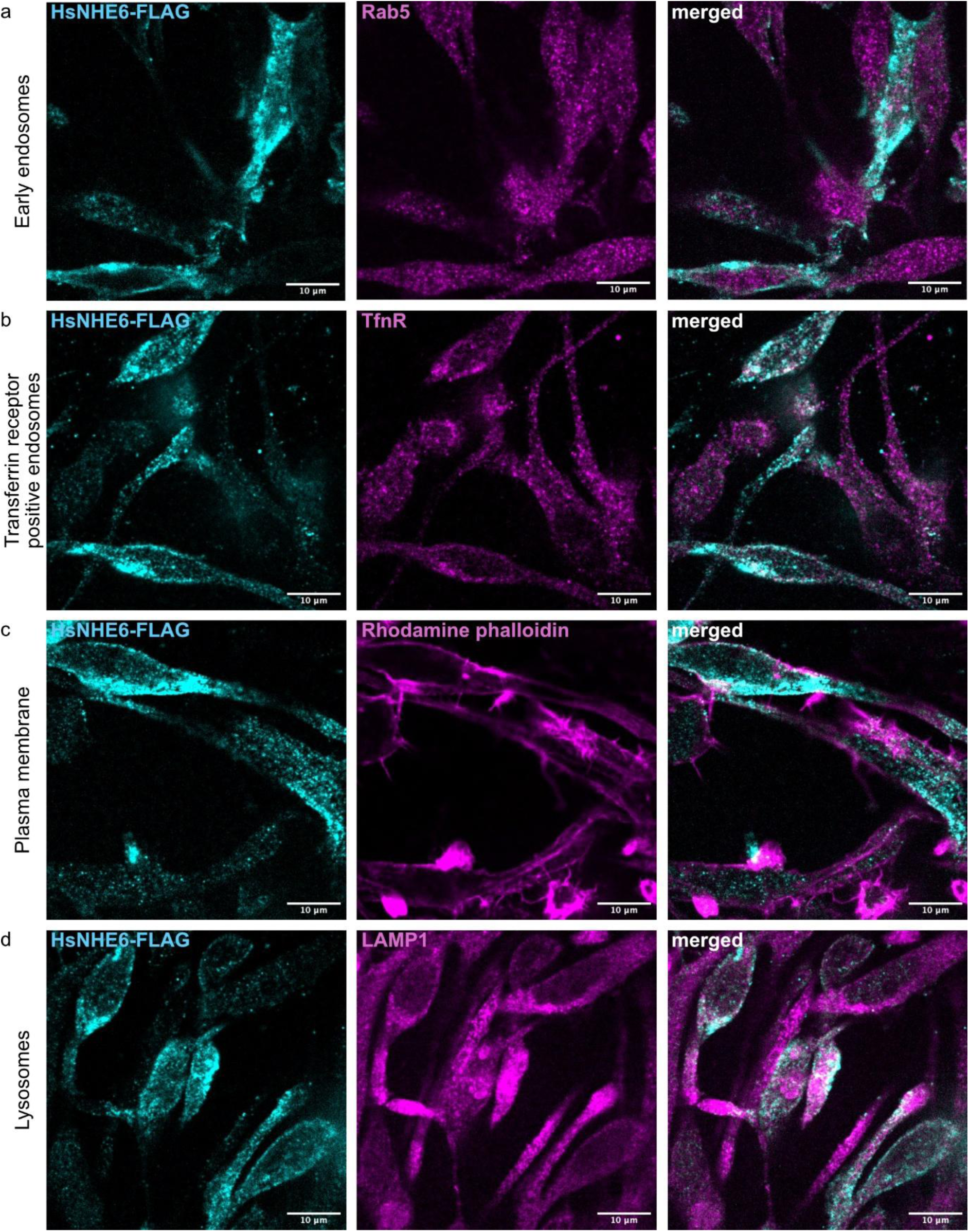
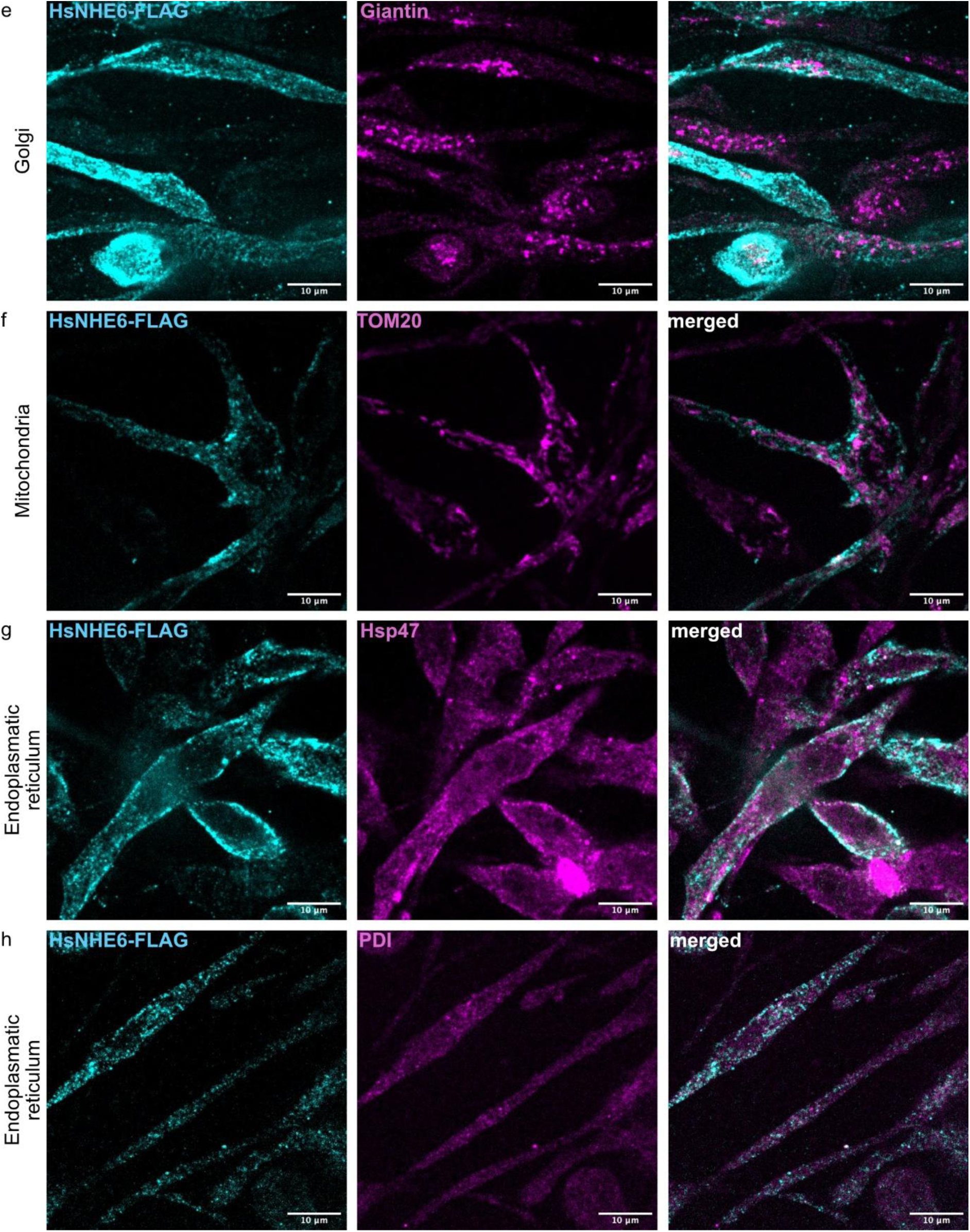
Colocalization of HsNHE6-Flag with cellular markers in FreeStyle293-F cells. Representative ICC colocalization images of HsNHE6-FLAG expression and cellular component markers. HsNHE6-FLAG in FreeStyle293-F partially colocalizes with **a,** early endosomal marker Rab5, **b,** Transferrin-Receptor (TfnR) positive endosomes, **c,** plasma membrane (F-actin, stained with Rhodamine phalloidin), and **d,** lysosomal marker LAMP-1, as well as with the **e,** Golgi protein Giantin. No colocalization was observed for **f,** mitochondria and **g, h,** ER. All scale bars correspond to 10 μm. Two independent biological experiments per condition.

**Supplementary Fig. 8.**
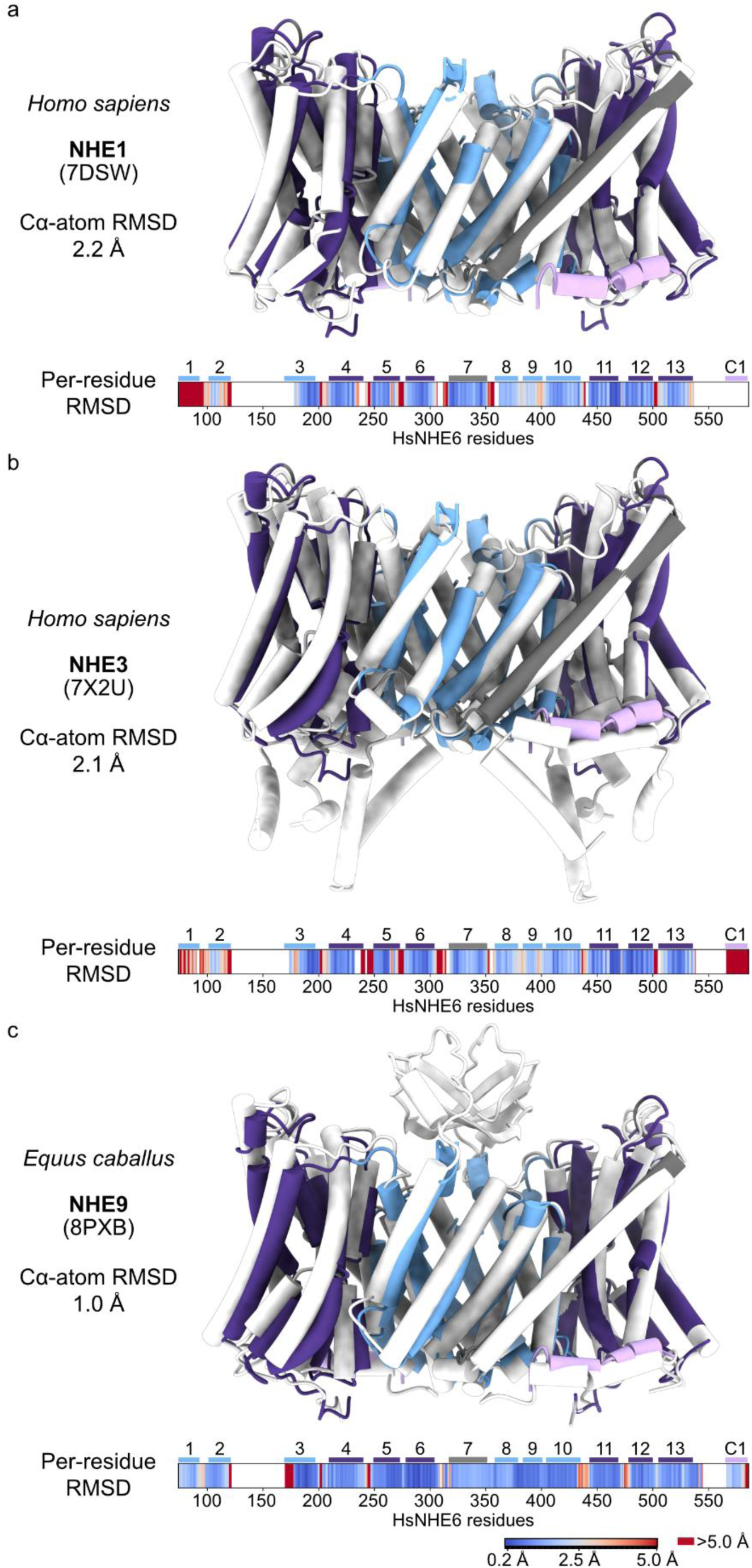
Comparison of HsNHE6 with published structural models of other NHEs. Overlay of the HsNHE6 model (colored) with the models (white) of **a,** HsNHE1 (PDB ID 7DSW)^6^, **b,** HsNHE3 (PDB ID 7X2U)^7^ and **c,** EcNHE9 (PDB ID 8PXB)^8^. The per-residues RMSD plots show the differences between the C^α^ atoms of one protomer, calculated after overlaying the dimerization domain TMDs of a single HsNHE6 protomer with those of the corresponding NHE protomer. Residue range and TM markers represent HsNHE6, white areas represent areas that were not matched or missing in (at least) one of the structural models.

**Supplementary Fig. 9.**
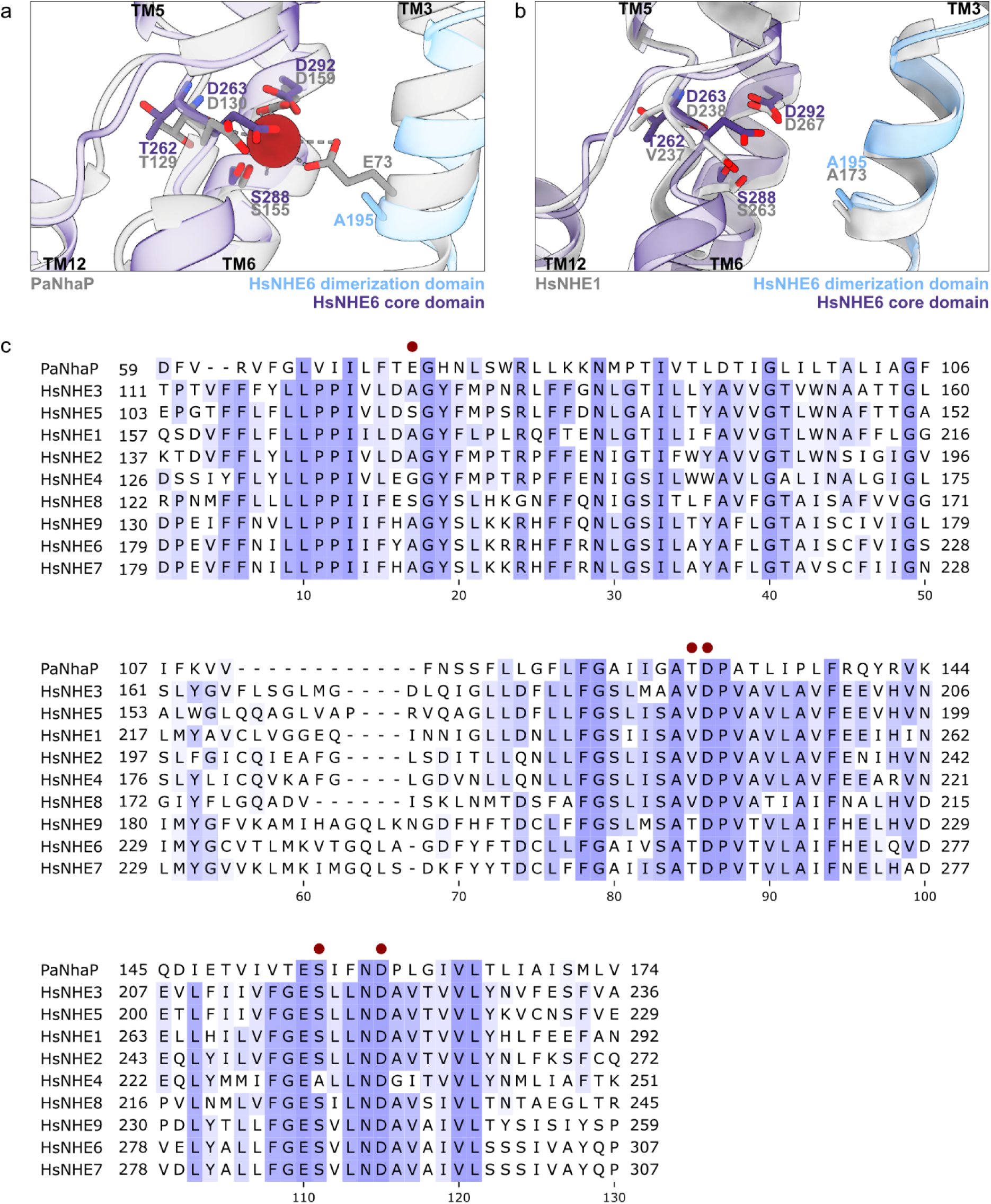
Comparison of the ion binding sites in HsNHE6, HsNHE1 and PaNhaP. **a,** Overlay of the structural models of HsNHE6 (violet and blue) with the structural model of PaNhaP (gray, PDB ID 4CZA)^9^ at the ion binding site. The side chains of the ion binding residues are shown as well as the thallium ion (red) of the PaNhaP structure. **b,** Overlay of the structural model of HsNHE6 (violet and blue) with the structural model of human NHE1 (PDB ID 7DSW)^6^ in the inward-facing conformation. The side chains of the ion binding residues are shown. **c,** Sequence alignment of human NHEs and PaNhaP covering the ion binding site. The red dots indicate residues directly involved in ion binding.

**Supplementary Fig. 10.**
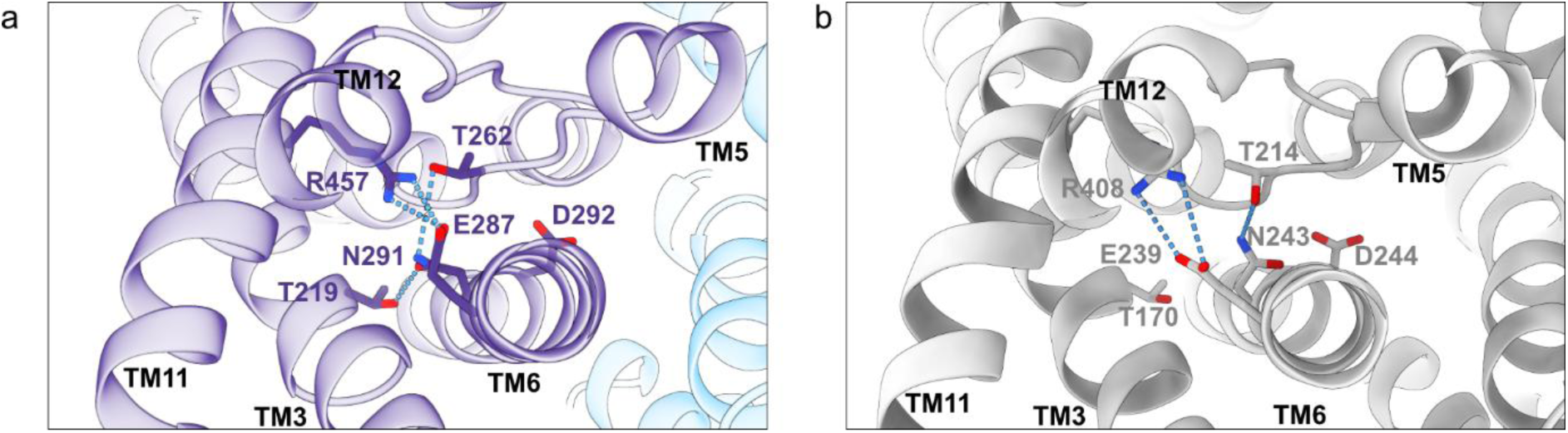
Stabilizing interactions in HsNHE6 and EcNHE9. **a,** Cartoon representation of HsNHE6, highlighting the salt bridge between E287^TM6^ and R457^TM11^ and the hydrogen bonds between N291^TM6^, T262^TM5^, and T219^TM3^. **b,** Cartoon representation of EcNHE9 (PDB ID 8PXB)^8^, highlighting the salt bridge between E239^TM6^ and R408^TM11^ and the hydrogen bonds between N242^TM6^, T214^TM5^, and T170^TM3^. The bonds are depicted in blue dashed lines.

**Supplementary Fig. 11.**
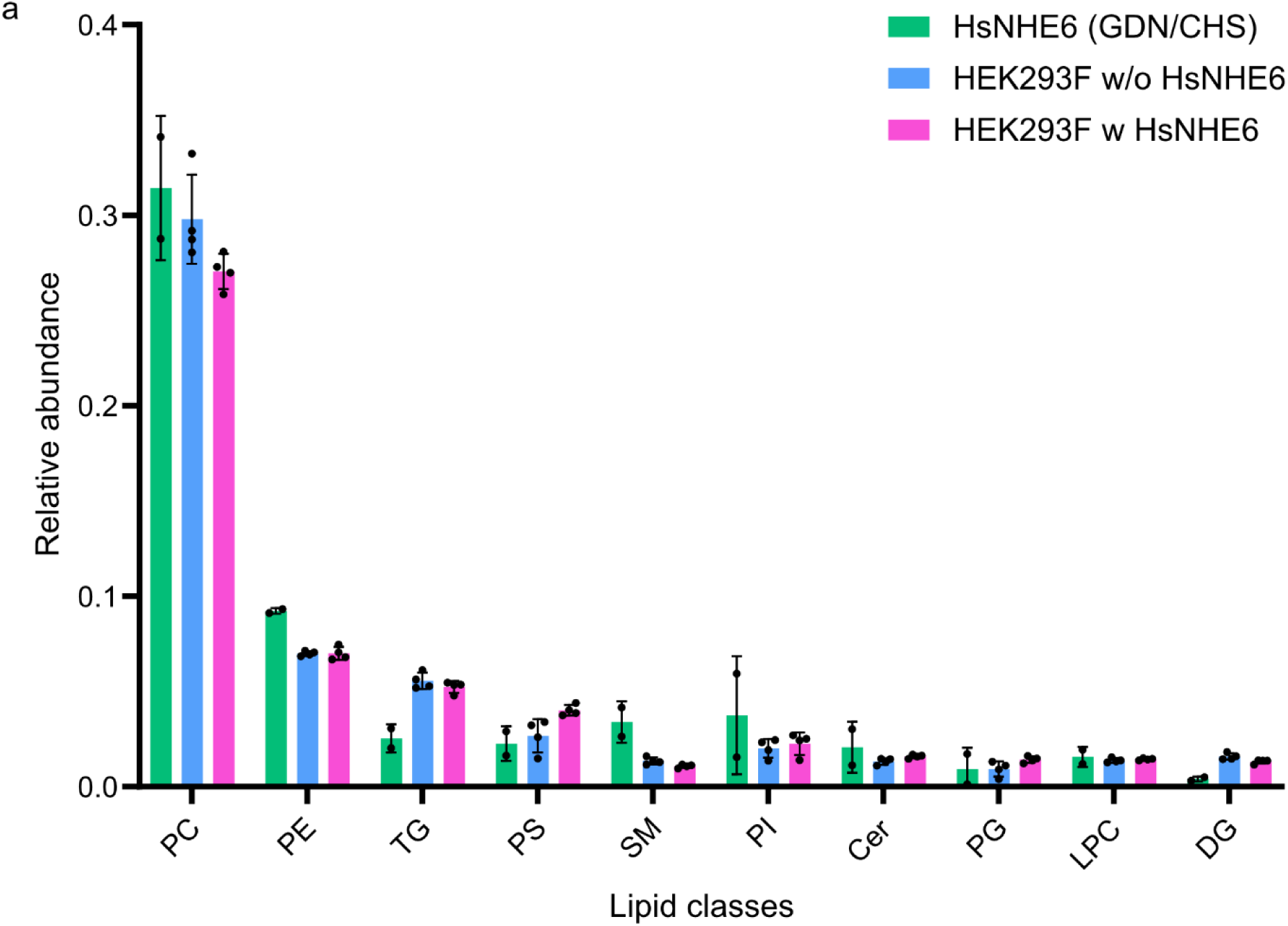
Lipid analysis of HsNHE6. **a,** Lipidomics analysis of detergent-purified HsNHE6 (green), detergent-solubilized HEK293F cells without overexpressed HsNHE6 (blue) and detergent-solubilized HEK293F cells with HsNHE6 expressed (magenta). The plot shows the top 10 lipid classes detected in the purified HsNHE6 sample. The signal of the individual identified lipid classes were computed relative to total raw signal including everything in the sample.

**Supplementary Fig. 12.**
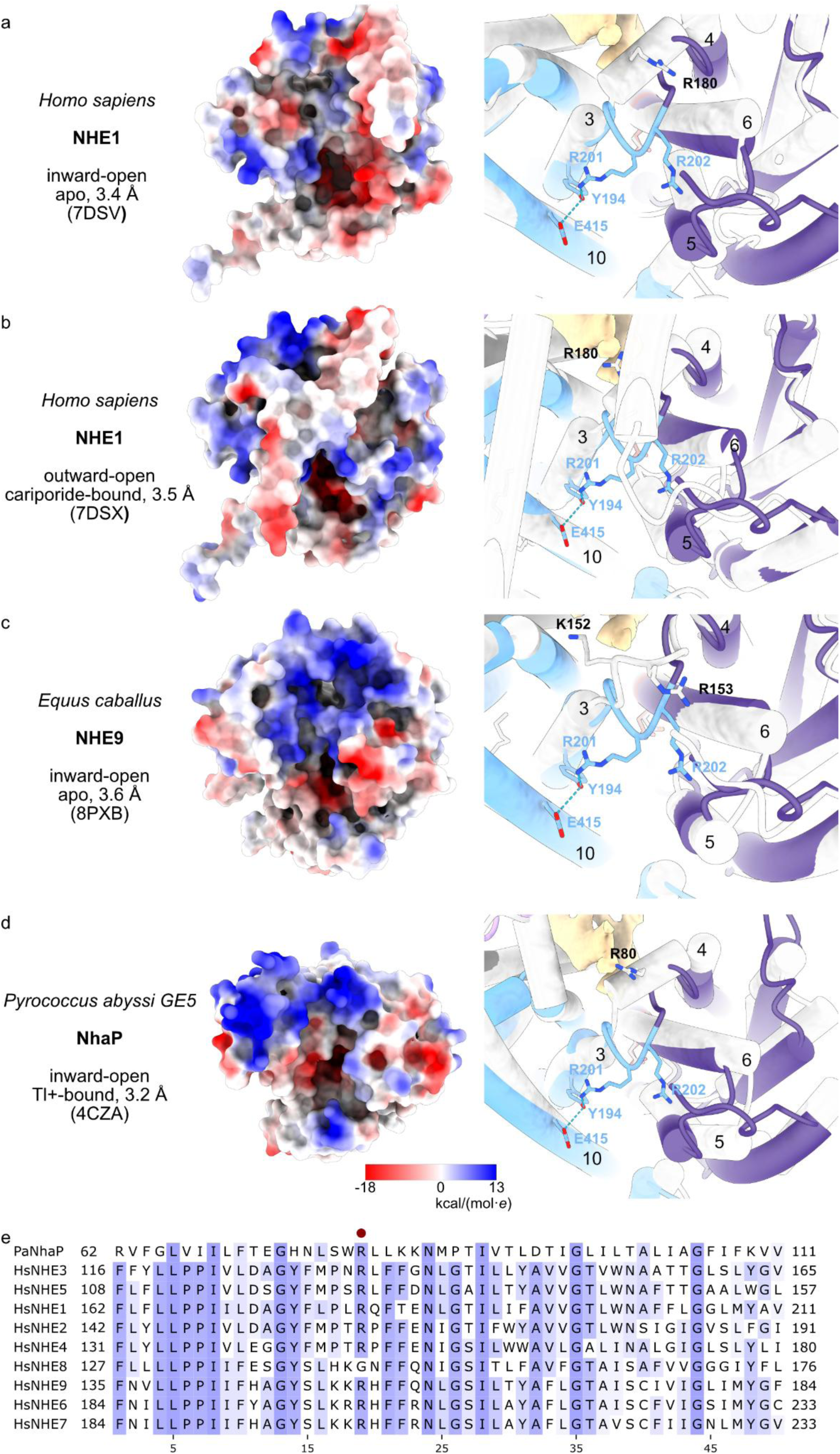
Comparison of the NHE ion-binding gates in HsNHE1, EcNHE9, PaNhaP with that in HsNHE6. Electrostatic surface representation (left) and cartoon representation (right) of one NHE protomer as seen from the cytoplasm. HsNHE6 model (blue and violet) overlayed with the respective NHE model (gray). **a,** HsNHE1 in inward-open conformation (PDB ID 7DSV)^6^, **b,** HsNHE1 in outward-open conformation (PDB ID 7DSX)^6^, **c,** EcNHE9 in inward-open conformation (PDB ID 8PXB)^8^ and **d,** PaNhaP in inward-open conformation (PDB ID 4CZA)^9^. **e,** Sequence alignment of SLC9A family members with PaNhaP, the highly conserved arginine between TM3 and -4 is marked (red dot).

**Supplementary Fig. 13.**
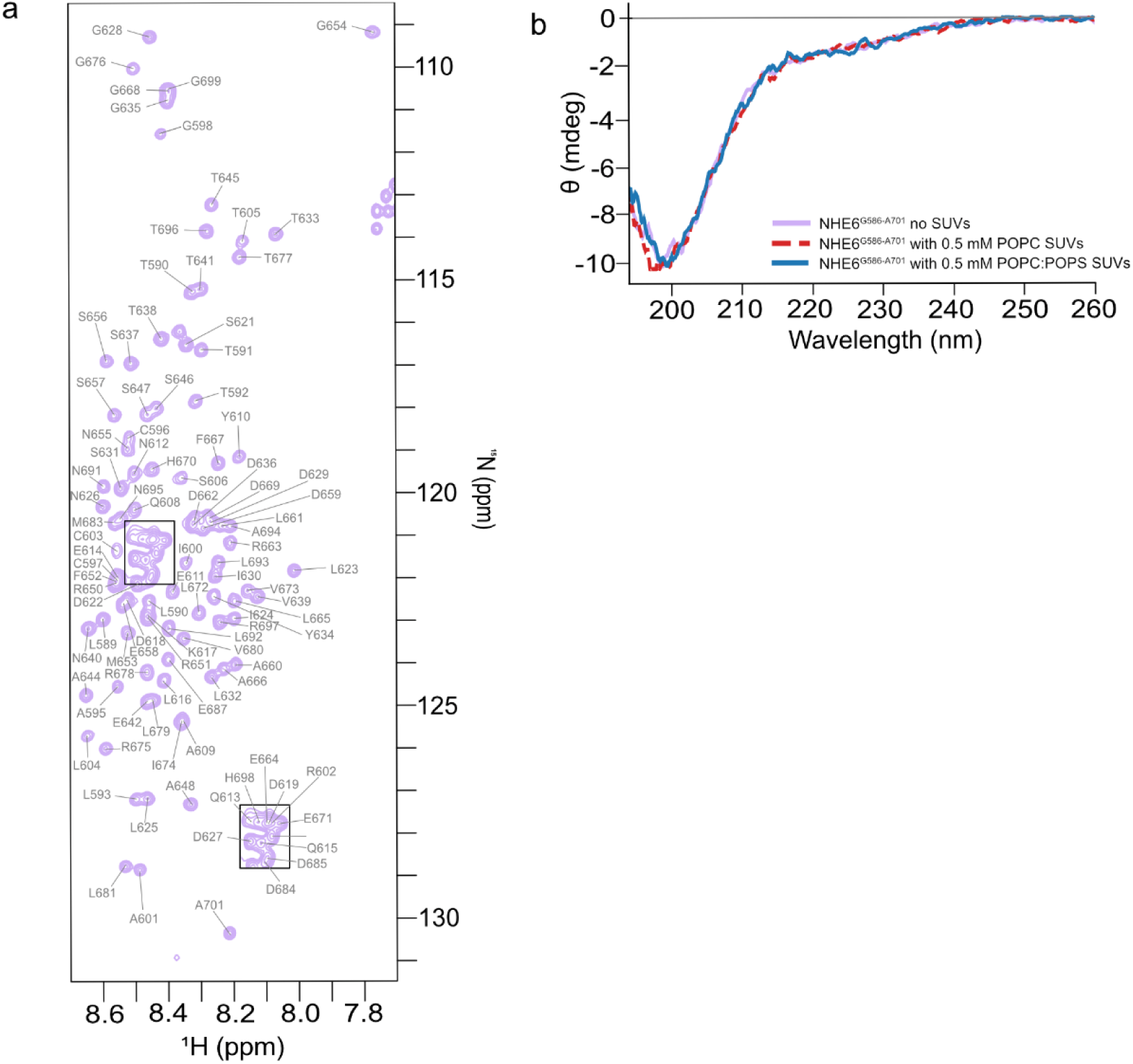
NMR and CD data for HsNHE6 C-terminus. **a,** ^1^H,^15^N HSQC spectrum of HsNHE6^G586-A701^ with assignments. **b,** Far-UV CD spectra of HsNHE6^G586-A701^ without SUVs (purple), and with either 0.5 mM POPC SUVs (dashed red) or 0.5 mM POPC:POPS (3:1) SUVs (blue).

**Supplementary Fig. 14.**
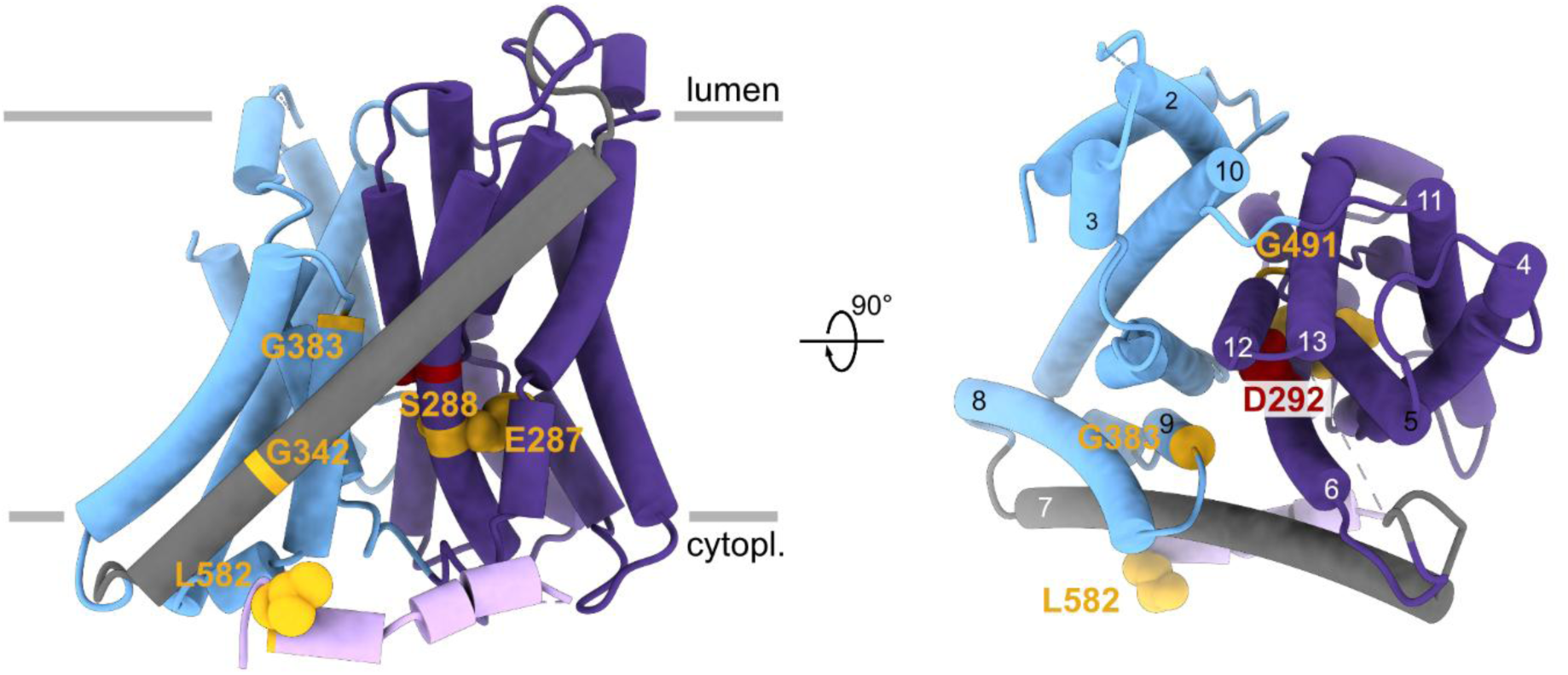
Christianson syndrome (CS)-causing missense mutations in HsNHE6. HsNHE6 monomer as vied in the membrane plane (left) and from the lumen (right) views, colored as in Fig 1a with the highly conserved D292 in red. The CS-causing missense mutations are colored in dark yellow. Amino acid side chains are shown as spheres.

### Supplementary Tables

**Supplementary Table 1.**
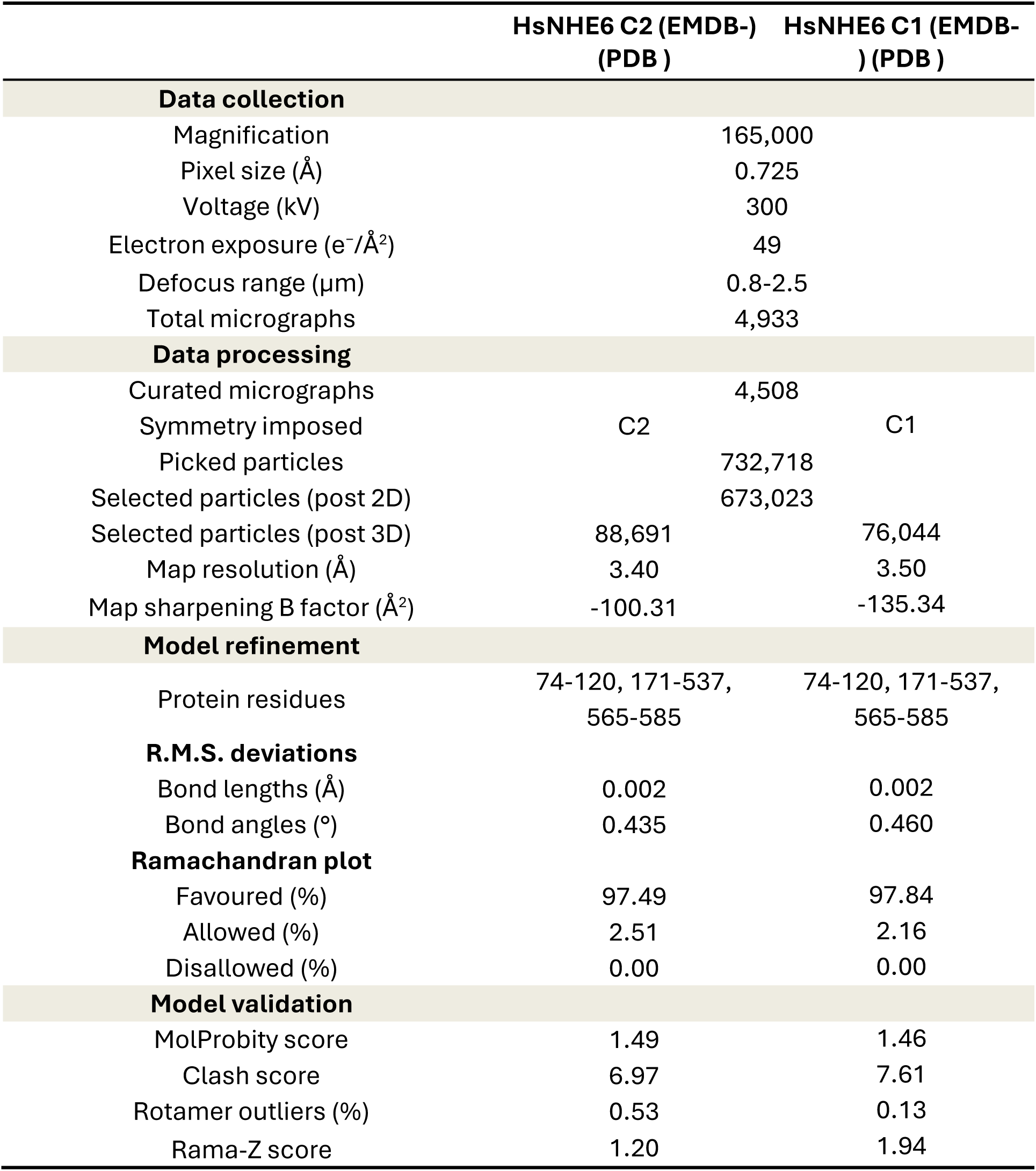
Data collection, processing and refinement statistics of HsNHE6 cryo-EM data.

**Supplementary Table 2.** Overview of Christianson syndrome-causing mutations in HsNHE6 based on Kavanaugh *et al.* and Ilie *et al.*^10,11^.

| Missense mutations |  | Nonsense mutations |  | Frameshift mutations |  |
| --- | --- | --- | --- | --- | --- |
| mutations | location | mutations | location | mutations | location |
| Δ287-288 | TM6 | R500* | TM12 | F183fs*1 | TM3 |
| G342R | TM7 | W523* | TM13 | I211Yfs*53 | TM4 |
| G383D | TM9 | S544* | C-terminus | T318fs*59 | TM7 |
| G491D | TM12 | W570* | C1 | H408Nfs*2 | TM10 |
| L582P | C1 |  |  | N460Kfs*12 | TM11 |
|  |  |  |  | L468Ifs*15 | TM11 |
|  |  |  |  | R472Kfs*4 | IL6 |

**Supplementary Table 3.**
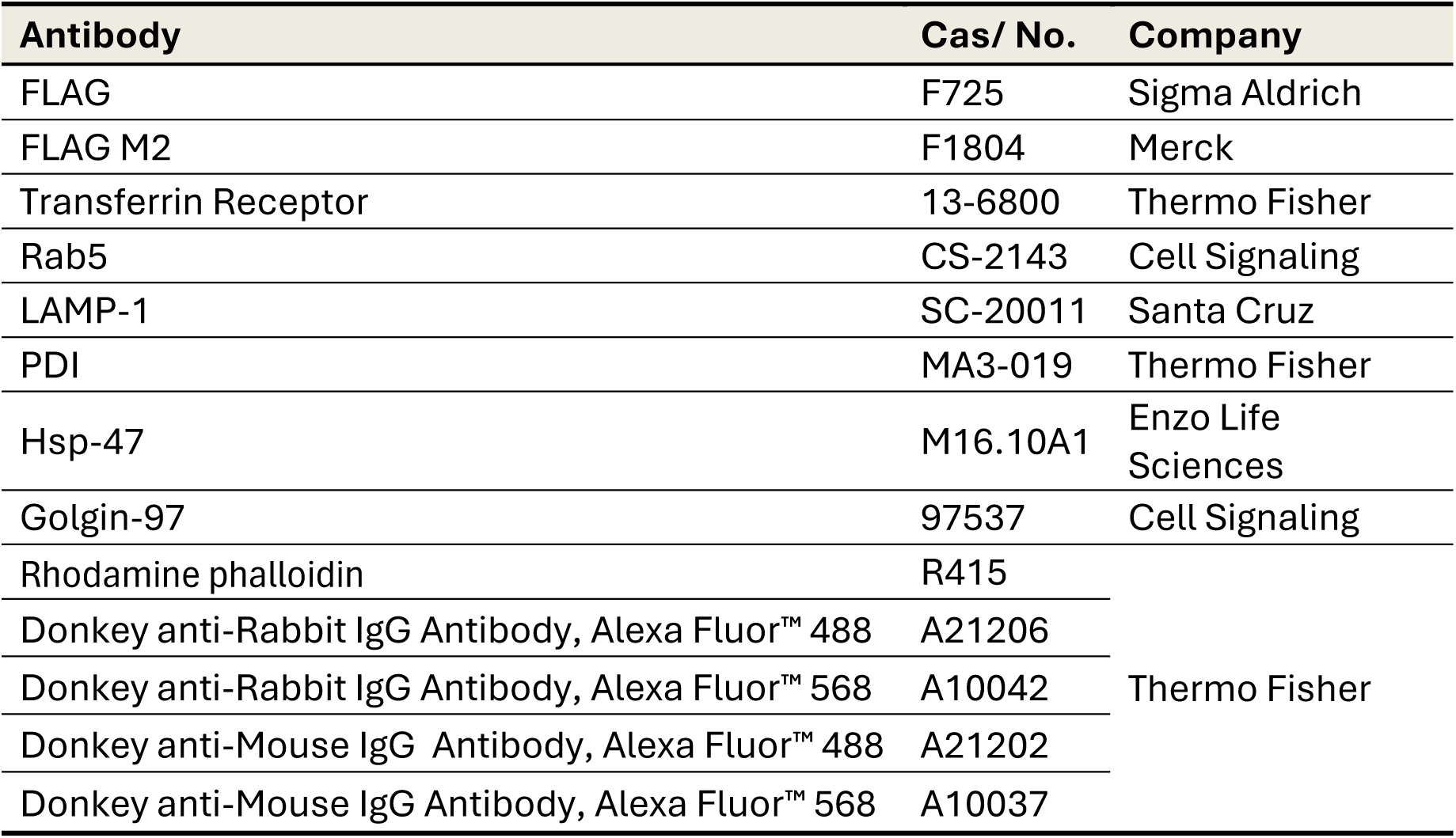
Primary and secondary antibody list for Co-localization experiment.

